# When do bursts matter in the motor cortex? Investigating changes in the intermittencies of beta rhythms associated with movement states

**DOI:** 10.1101/2022.06.22.497199

**Authors:** Timothy O. West, Benoit Duchet, Simon F. Farmer, Karl J. Friston, Hayriye Cagnan

## Abstract

Time series of brain activity recorded from different anatomical regions and in different behavioural states and pathologies can be summarised by the power spectrum. Recently, attention has shifted to characterising the properties of changing temporal dynamics in rhythmic neural activity. Here, we present evidence from electrocorticography recordings made from the motor cortex to show that, dependent on the specific motor context, the statistics of temporal transients in beta frequency (14-30 Hz) rhythms (i.e., bursts) can significantly add to the description of states such rest, movement preparation, movement execution, and movement imagery. We show that the statistics of burst duration and amplitude can significantly improve the classification of motor states and that burst features reflect nonlinearities not detectable in the power spectrum, with states increasing in order of nonlinearity from movement execution to movement preparation to rest. Further, we provide mechanistic explanations for these features by fitting models of the motor cortical microcircuit to the empirical data and investigate how dynamical instabilities interact with noise to generate burst dynamics. Finally, we examine how beta bursting in motor cortex may influence the integration of exogenous inputs to the cortex and suggest that properties of spontaneous activity cannot be reliably used to infer the response of the cortex to external inputs. These findings have significance for the classification of motor states, for instance in novel brain-computer interfaces. Critically, we increase the understanding of how transient brain rhythms may contribute to cortical processing, which in turn, may inform novel approaches for its modulation with brain stimulation.

## 1 Introduction

Rhythmic activity from populations of neurons, as is routinely summarised in the power spectrum, is often taken to be sufficient to characterise neural activity from different brain regions (Keitel and Gross 2016; Mahjoory et al. 2020), behavioural states (Siegel et al. 2012), and pathologies (Brown et al. 2001; Schnitzler and Gross 2005). However, when analysed in time, neural rhythms often resolve into a succession of intermittent, transient events (Baker et al. 2014; van Ede et al. 2018; Fingelkurts and Fingelkurts 2010; Freeman 2004; Friston 1997) that can appear as sustained oscillations when investigated using trial averaged analyses (van Ede et al. 2018; Jones 2016). To understand how alterations in power are underwritten by the temporal dynamics of neural rhythms, it is necessary to explicitly quantify the duration, amplitude, and rate of transient events (Heideman et al. 2020).

Temporal intermittencies in neural rhythms (i.e., “bursts”) are known to be important in behaviours such as sleep (Adamantidis et al. 2019) and working memory (Lundqvist et al. 2016). In the healthy motor system, changes in the temporal patterning of beta frequency (14-30 Hz) activity can predict behaviour beyond that achieved when using just the amplitude of beta activity (Enz et al. 2021; Hannah et al. 2020; Shin et al. 2017; Wessel 2020). Further, beta burst dynamics appear to be significantly altered in Parkinsonism (Cagnan et al. 2019; Deffains et al. 2018; Tinkhauser et al. 2017b), where they form a major target for adaptive deep brain stimulation (Little et al. 2016; Tinkhauser et al. 2017a). An important consideration for therapeutic stimulation specificity is discriminating between pathological and healthy motor activity. Properties of transient activity can, in principle, improve classification accuracy and thus increase the specificity of stimulation effects.

In the context of motor behaviour, preparation and execution have been conventionally described in terms of event related synchronization and desynchronization in the beta frequency band (Pfurtscheller and Lopes da Silva 1999). Movement imagery has also been linked to event related desynchronization albeit with less power decrease in beta when compared to movement execution (Pfurtscheller and Neuper 1997). When temporally resolved, changes in the rate and timing of beta bursts are associated with movement preparation, planning, termination or cancellation (Diesburg et al. 2021; Feingold et al. 2015; Khanna and Carmena 2017; Little et al. 2019; Torrecillos et al. 2018; Tzagarakis et al. 2010; Wessel 2020). Additionally, the occurrence of beta bursts is associated with effects that persist beyond their termination (Khanna and Carmena 2017; Torrecillos et al. 2018). It has been suggested that bursts reflect a competition between endogenous processing and external sensory responses that bias perception in the cortex (Karvat et al. 2021).

Taken together, we hypothesize that (1) the temporal properties of beta bursts are altered between different movement states; (2) these changes in dynamics reflect altered responses of the motor cortex to stochastic inputs, that arise from a reconfiguration of the underlying microcircuit, and thus (3) bursts reflect a rebalancing of how the cortex integrates between spontaneous and exogenous inputs.

To date, the mechanisms underlying burst activity have been described using relatively simple models, such as an excitatory/inhibitory network of Wilson-Cowan populations (Duchet et al. 2021; Powanwe and Longtin 2019; Xing et al. 2012) that are motivated by pyramidal-interneuron models of beta generation (Jensen et al. 2005; Kopell et al. 2011). These studies indicate that burst statistics are determined by interactions between synaptic noise and the connectivity parameters of any given model. This suggests that models constrained using burst statistics can more accurately infer underlying connectivity across states, particularly in more complex models of the motor cortex. In models incorporating a more complete structure, previous work has demonstrated the importance of laminar specific corticothalamic inputs, which given the right timing can generate short, high amplitude beta events (Sherman et al. 2016). Whilst these models have been useful in understanding how to either experimentally or therapeutically modulate the mechanisms that give rise to beta bursts, it is still not known how changes in burst statistics during different stages of movement are underwritten by alterations in cortical microcircuitry.

This present work aims to establish how alterations of the cortical microcircuitry during motor behaviour are manifest in the burst statistics of beta rhythms recorded from large scale neuronal activity. To this end, we use a library of publicly available electrocorticography (ECoG) data recorded from participants performing a range of motor tasks (Miller 2019). We first investigated how rhythmic burst features in these data may enhance the classification of different motor stages—such as movement preparation, execution, and imagery—by providing information beyond that available in the time averaged spectra. Secondly, using computational models of the motor cortex microcircuit fitted to the burst statistics and spectra of the ECoG data, we characterise how biophysical parameters may modulate bursting dynamics in different brain states and investigate whether the changes in the expression of beta bursts can reflect the altering balance between spontaneous and exogenous drives to the motor cortex.

## 2 Methods

### 2.1 Electrocorticography and Experimental Recordings

All experimental data was taken from an openly available library (Miller 2019) published for use without restriction (https://searchworks.stanford.edu/view/zk881ps0522). Recordings were made for anatomical mapping in patients with epilepsy at Harborview Hospital, Seattle, WA, USA. All patients provided informed written consent, under experimental protocols approved by the Institutional Review Board of the University of Washington (see supplementary information 1). Data were recorded at the bedside using Synamps2 amplifiers (Compumedics Neuroscan). Visual stimuli were presented using a monitor running BCI2000 stimulus and acquisition programs (Schalk et al. 2004) that were also synchronized to behavioural feedback in the tasks (see below). Electrocorticography was recorded using grids and/or strips of platinum subdural electrodes placed via craniotomy. Electrodes had a 4 mm diameter (2.3 mm exposed), 1 cm interelectrode distance and embedded in silastic (figure 1B). Electrical potentials were recorded at 1 KHz using a scalp/mastoid reference and ground. Hardware imposed a bandpass filter from 0.15 to 200 Hz. Locations of electrodes were confirmed using post-operative radiography. Exact details of the electrode localization methods can be found in Miller (2019).

**Figure 1.**
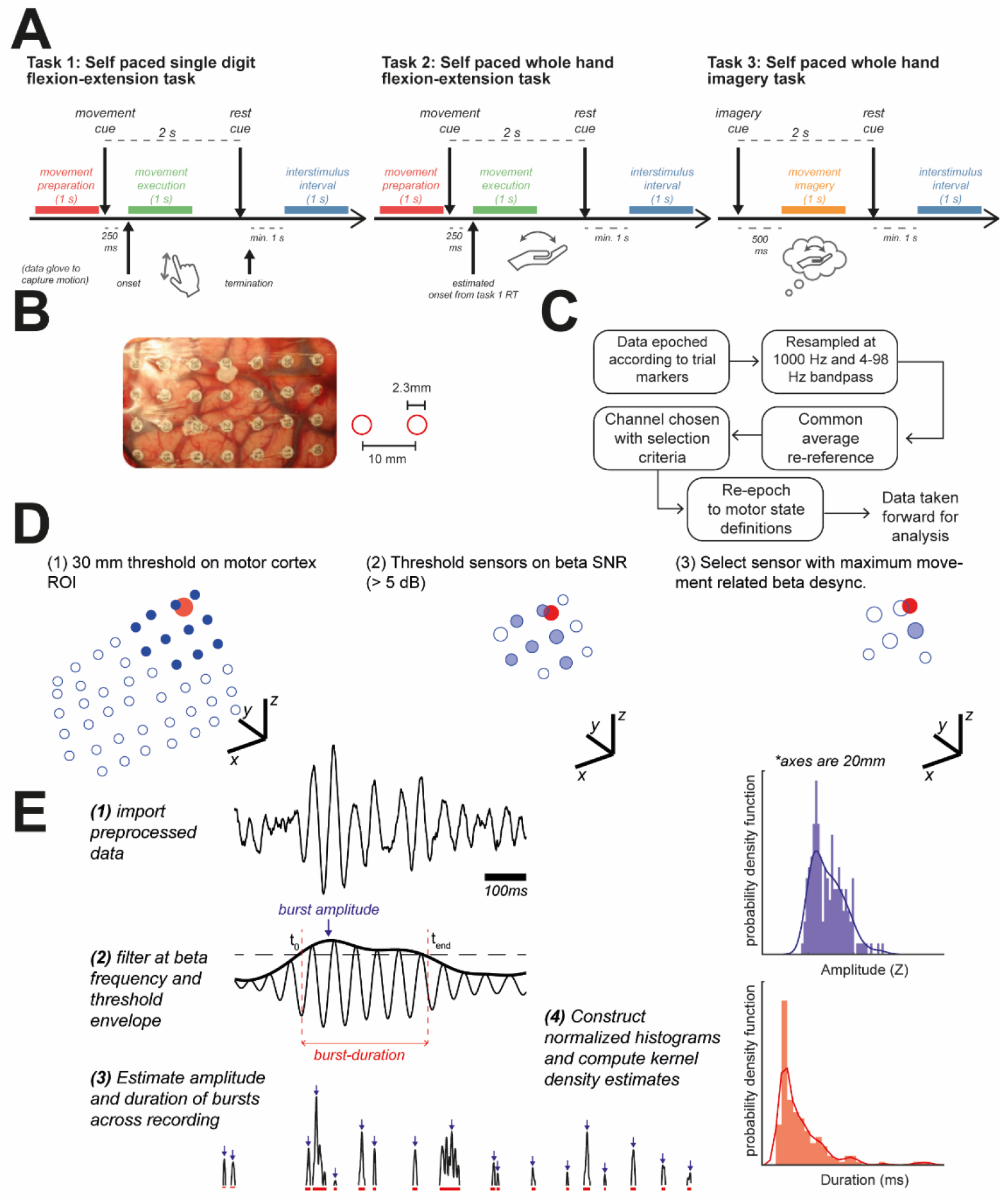
Illustrated criteria for selection of ECoG channels and computed data features: spectra, and distributions of burst amplitudes and durations. **(A)** Data was taken from three motor tasks, requiring either self-paced flexion/extension of individual digits (task 1); or flexion/extension of whole hand (task 2); or imagery of whole hand movement (task 3). Data was epoched according to timings relative to given in figure. **(B)** Electrocorticography was recorded with grids of platinum electrodes placed subdurally via craniotomy. Inset schematics give scale: electrodes had a 2.3 mm diameter with 10 mm spacing. **(C)** Procedures for preprocessing data. **(D)** Illustration of channel selection procedure. Candidate ECoG channels (blue open circles) were selected (filled blue circles) using a 30 mm search radius of the ROI (MNI coordinate: [±37 −25 62]; red circle). All channels were thresholded at a 5 dB SNR threshold for the beak peak (see methods), finally channels were selected using the maximum movement related beta desynchronization. **(E)** Illustration of envelope threshold procedure to identify bursts. Samples of burst amplitudes and durations were used to construct histograms. The summaries of these distributions were then taken as the kernel estimate to the probability density function. Image of electrocorticogram in panel (B) is reprinted by permission from Springer Nature, Nature Human Behaviour (Miller 2019).

**Figure 2.**
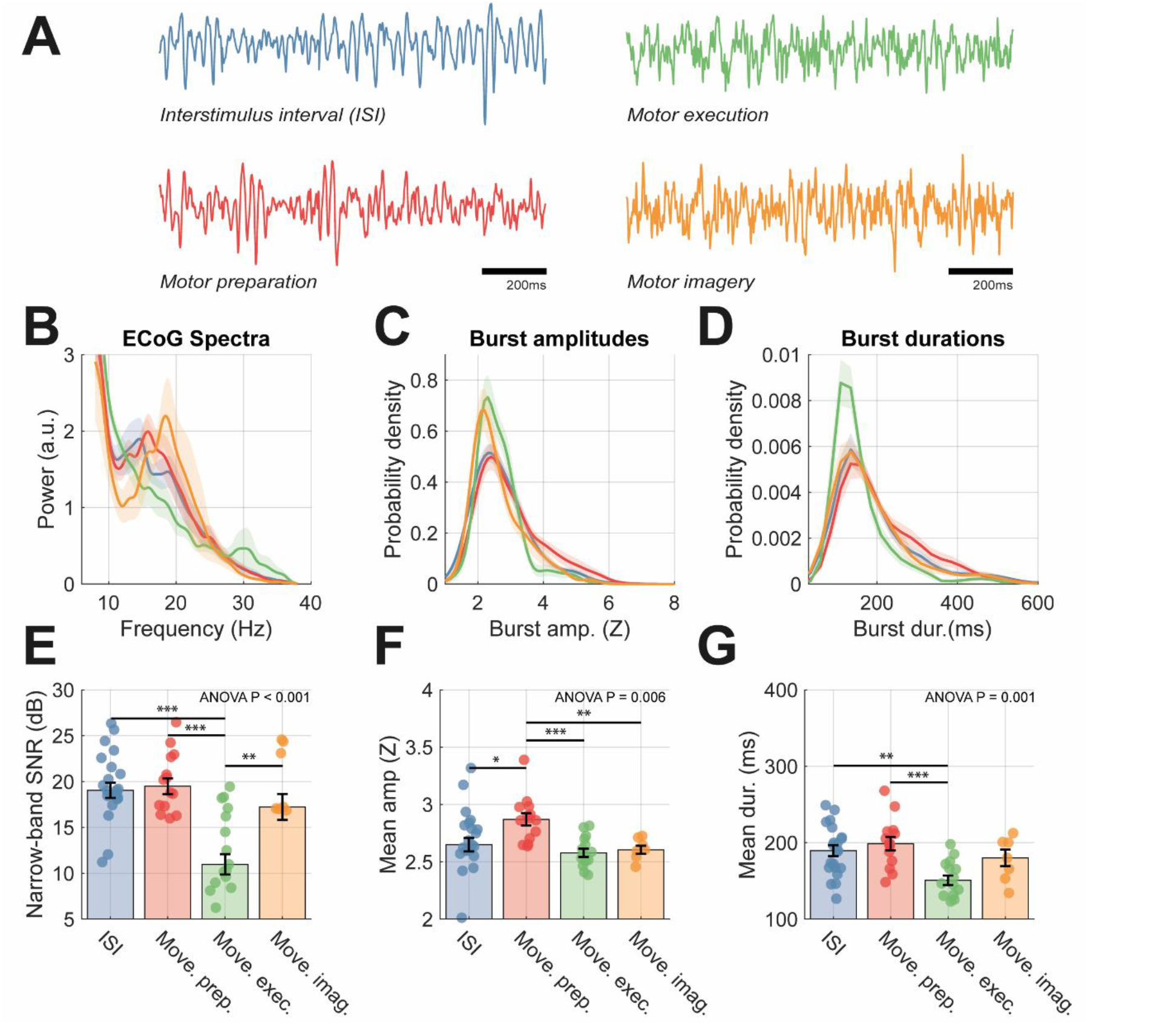
Analysis of recordings from selected ECoG sensors exhibit changes in properties of both spectral and burst features between motor states. Analyses were split among motor states: interstimulus interval (blue), movement preparation (red), movement execution (green), and motor imagery (orange). **(A)** Example 2 second time series of ECoG recordings for different motor states. Clear bursts of beta activity are apparent in ISI, movement preparation, and imagery states. **(B)** Group average of normalized power spectra, **(C)** probability density of burst amplitudes (given as Z scores), and **(D)** probability density of burst durations (ms). Bar plots in (E-G) show data from individuals overlaid, with mean and standard distributions indicated by error bars. Data is shown for: **(E)** narrow-band SNR (dB); **(F) (F)** median burst duration (ms); **(G)** mean burst amplitude (Z score). Statistics indicate results of one-way ANOVA with bars indicating respective significant post-hoc t-tests between pairs of states. An analysis of the predictive value of burst vs spectral features in classifying motor states can be found in supplementary figure 2.

Data were taken from three different tasks as summarised below. For details of task structure and trial definitions please see figure 1A. Subject numbers represent the initial total available for each task, some subjects participated in more than one task. Data selection procedures are given in section 2.2.

*Dataset 1: Self-Paced Finger Movements (n = 9)* – originally reported in Miller et al. (2012). Participants were cued with a word displayed on a bedside monitor indicating which digit to perform a self-paced flexion and extension during a 2 s movement trial. Trials typically comprise 2-5 movements as recorded using a data glove. Movement blocks were interleaved with 2 s rest trials.

*Dataset 2: Basic Motor (n = 19) –* originally reported in Miller et al. (2007b) and Miller et al. (2010). Participants were asked to make either a simple repetitive flexion and extension of all the fingers, or a protrusion and retraction of the tongue at self-paced rate (∼2 Hz). Patients were cued with a picture of the body part to move, presented on a screen.

*Dataset 3: Motor Imagery (n = 7) –* originally reported in Miller et al. (2010). Participants were asked to imagine making a simple repetitive flexion and extension of the fingers, or protrusion/protraction of the tongue at a self-paced rate (∼2 Hz), matched to the task described for dataset 2. Imagery was intended to be kinaesthetic rather than visual-i.e., “imagine making the motion, not what it looked like”. Movement blocks lasted 2 or 3 s and were always followed by rest intervals of the same length.

### 2.2 Pre-processing and Criteria for Data Selection

All ECoG recordings were processed as summarised in figure 1C. Large scale artefacts common across sensors were reduced by referencing electrodes to the common average. Channels with significant artefacts or epileptiform activity were visually rejected and excluded from the common average. Finally, data were filtered between 4-98 Hz using a zero-phase (i.e., forward-backward) FIR filter with −60 dB stopband attenuation. A 4 Hz passband was chosen to remove the influence of lower frequency rhythms in the SNR calculations for the beta peak. Data from each task were segmented to 1 second epochs. For each set of recordings, we selected one ECoG channel to carry forward for analysis. Data were selected to identify signals which were relevant to motor cortical activity (i.e., spatially close to primary motor cortex), of sufficient quality (i.e., good signal-to-noise of beta frequency activity), and functionally relevant (i.e., showing task related changes in synchrony). An illustration of the selection process can be seen in figure 1D. Channels were selected based on the following criteria: (1) select channels within 30mm of left or right primary motor cortex (MNI: [±37 −25 62]; Jha et al. (2015)); (2) threshold channels at +5 dB SNR for the beta band (14-30 Hz); (3) select channel based on maximum SNR change between rest and movement/imagery. If no channels were found that matched these criteria the subject was removed from further analysis. The number of subjects whose data was carried forward for further analysis was: 5/9 subjects from dataset 1; 10/19 subjects from dataset 2; and 4/7 from dataset 3.

Details of epoching are illustrated in figure 1A. For dataset (1), kinematic data was available from a data glove worn during the experiment, and thus data was epoched according to movement onset (finger movements) determined using a threshold crossing on the smoothed movement traces. Data was segmented into *movement preparation* (−1250 ms to −250 ms relative to movement onset) and *movement execution* (0 ms to +1000 ms relative to movement onset) and then 1 s *interstimulus intervals* (ISI) blocks taken in between movement cues. ISI blocks were always at least 1 s away from a movement cue or movement termination. Note that we left a 250 ms gap prior to movement onset, as we wanted to avoid non-stationarities while beta exhibited movement related desynchronization. For datasets 2 and 3, movement kinematics were not available, and movement or imagination was cued by on-screen instructions. We therefore estimated movement onset using reaction times from dataset 1. If a subject also participated in dataset 1, we used their median subject-specific reaction time. For all other subjects, we used the group median. We took blocks of *movement execution* and *movement imagery* starting cue onset + reaction time (lasting for 1 s). Movement preparation was defined as before.

### 2.3 Data Features: Spectra and Distributions of Burst Amplitude/Duration

Time series data were summarised using features derived from both spectra and bursts. We computed power spectral densities using Welch’s periodogram method with no overlap and a 1 s Hanning window. Spectra were summarised using their peak frequency, wide-band SNR, and narrow-band SNR within the beta band (14-30 Hz) (see supplementary methods I). Spectra used for model fitting were pre-processed to remove the 1/f aperiodic background so as to isolate peaks at beta frequency only (see figure 3A and supplementary methods II).

**Figure 3.**
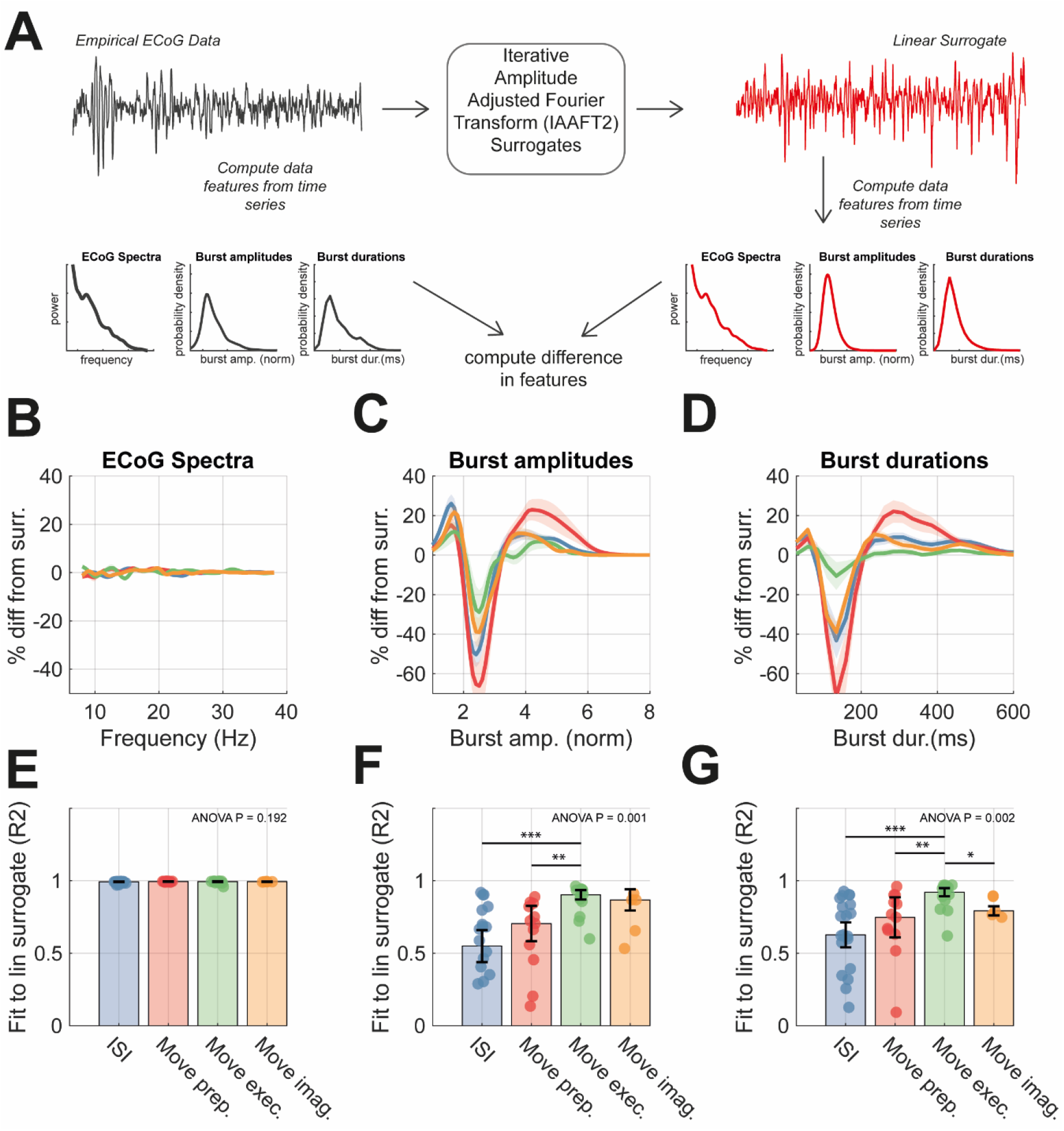
Comparison of empirical ECoG data with linear surrogates show that burst features represent significant signal nonlinearity that is modulated across conditions. **(A)** The Iterative amplitude adjusted Fourier transform (IAFFT; see methods) was used to construct spectra-matched, linear surrogates (right) for each of the ECoG recordings (left). Spectral and burst features were computed for each signal, and the difference between the surrogate and empirical features were compared to assess the extent to which nonlinearities were present in data from the four motor states. **(B)** Plots showing the averaged difference between surrogate and empirical power spectra (computed as a percentage change). **(C)** Same as (B) but for distributions of burst amplitudes. **(D)** Same as (B) but for burst duration distributions. **(E)** Bar chart indicating the median goodness-of-fit of the surrogate to the empirical data feature with IQR shown by error bars. **(F)** Same as (E) but for burst amplitude distributions. **(G)** Same as (E) but for burst duration distributions. Statistics indicate results of one-way ANOVA with bars indicating respective significant post-hoc t-tests between pairs of states.

Bursts were defined using a threshold on the bandlimited envelope (Cagnan et al. 2019; Tinkhauser et al. 2017b). Note that thresholds were epoch specific (local) to avoid the bias towards burst effects reflecting simple differences in signal to noise that can occur with a common threshold (Schmidt et al. 2020). For an illustration of burst definitions and the formation of summary statistics of burst properties, see figure 1E. Details of the procedure are given in supplementary methods III.

Overall, spectral features comprised: (1) wide-band SNR, (2) narrow-band SNR, (3) peak frequency. Burst features comprised: (5 and 6) mean and standard deviation of burst duration; (7 and 8) mean and standard deviation of burst amplitude; (9 and 10) mean and standard deviation of the inter-burst intervals. Statistical tests were computed on log transformed data. For all features except peak frequency, a one-way ANOVA and post-hoc t-tests were used to test for changes in means of features between motor states. The distribution of peak frequencies was not found to be normal, therefore, a Kruskal-Wallis test plus post-hoc rank-sum tests were used to determine changes in mean.

### 2.4 Assessing Feature Nonlinearity: Comparison with Linear Surrogate Data

To assess the extent to which statistics of burst features in cortical signals encode information beyond that contained in the power spectrum—a data feature sufficient for linear systems—we used a comparison to surrogate data (Theiler et al. 1992). Following previous work characterising the degree of nonlinearity in beta bursts (Duchet et al. 2021), we adopt the use of Iterative Amplitude-Adjusted Fourier Transforms (IAAFT; Schreiber and Schmitz 1996). IAAFT surrogates method improves upon the simpler technique of constructing randomized-phase Fourier surrogates, by not only ensuring the power spectrum is preserved, but also that the signal’s probability density is preserved. This ensures that the surrogate reproduces the linear features of the data whilst destroying potential nonlinearities in the original time series. To compare data with IAFFT surrogates, we constructed 25 surrogate time series for each data set, and then took the feature average, computed in the same way as for the reference (i.e., the empirical or simulated) signals. We then computed the goodness-of-fit in terms of the R^2^, with R^2^ << 1 indicating significant deviation of a data feature from that expected in the equivalent linear process.

### 2.5 Classification of Functional States with a Support Vector Machine

To determine the ability of different data features to decode the functional state from neural activity we employed a classification approach. Prior to classification, we applied Linear Discriminant Analysis (LDA) to the data to reduce the dimensionality of the feature space to two LDA components. We then used a multiclass support vector machine (SVM) using error-correcting output codes (ECOC) to combine binary classifiers into an ensemble and applied this to the LDA feature space. Learners were implemented in MATLAB using iteratively optimized hyperparameters, and a Gaussian kernel set. Model performance was evaluated using five-fold cross validation and the area under the curve (AUC) of receiver operating characteristics (ROCs) across the folds. Plots of SVM decision bounds were computed using posterior probabilities of model predictions applied in a grid search across the feature space. In effect, these measures of classification accuracy constitute an empirical estimate of model evidence or marginal likelihood, where the model in question maps from a functional (motor) state to various data features.

### 2.6 Fitting a Model of Motor Cortex Population Activity

We used a neural mass model of population activity in the motor cortex microcircuit (i.e., Bhatt et al. (2016)). This neural (state space) model formulation follows from the Wilson-Cowan firing rate model (Vogels et al. 2005), and has been used previously to describe dynamics of beta oscillations in the cortico-basal ganglia circuit (Oswal et al. 2021; Pavlides et al. 2015). This model delivers the average firing rate in response to input currents generated by spike trains from connected populations. Interlaminar projections were modelled using a delayed connectivity matrix reflecting the pattern of connectivity outlined in figure 4A. The model is driven using 1/f^α^ noise generated using a fractional Gaussian process (Dietrich and Newsam 1993), with *α* a free parameter to be fit. For a full description of the model equations please see the supplementary methods IV. The model comprises three pyramidal cell layers (superficial *SP*, middle *MP*, and deep *DP*) plus one population of inhibitory interneurons (II). Each cell layer receives a self-inhibitory connection reflecting local synaptic gain control. The output of the model is a weighted sum (i.e., a lead field) of the layer specific firing rates with 80% contribution from deep layers, and 10% from superficial and middle.

**Figure 4.**
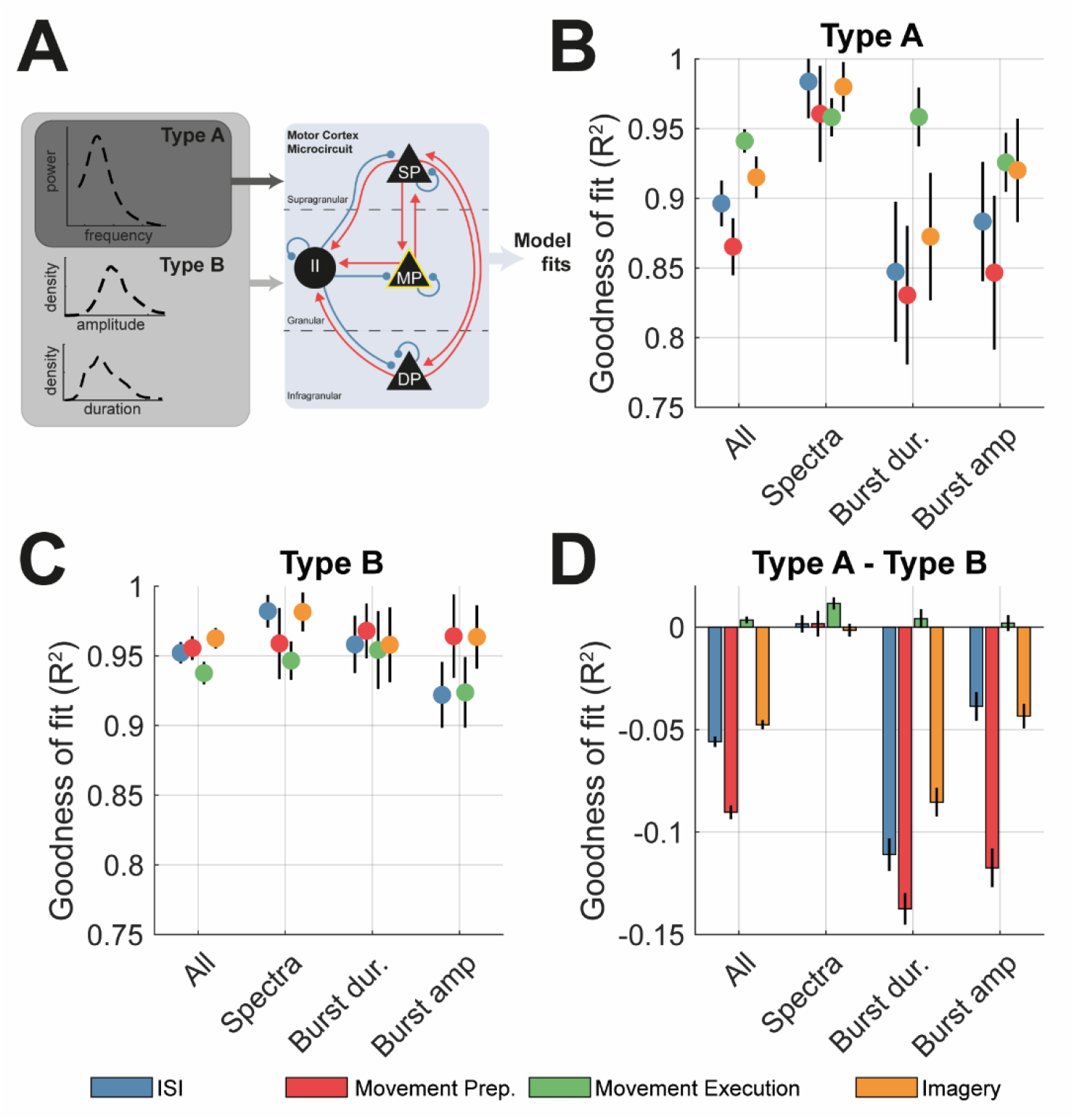
Comparison between type A (spectra only) and type B (spectra + burst features) fits of the motor cortex microcircuit demonstrates that spectral features are not sufficient to accurately constrain simulated burst parameters. Data features were constructed by simulating data using draws from the posterior distributions over parameters (n = 256). **(A)** Schematic of the motor cortex microcircuit model. Each black node represents a neural mass that is coupled with either excitatory (red) or inhibitory connections (blue). There are three pyramidal cell layers: superficial (SP), middle (MP), and deep (DP), plus an inhibitory interneuron (II) population. Model parameters were constrained using either pre-processed spectra (type A) or both spectra and burst features (type B) **(B)** Summary of the median ±SEM goodness of fit (R^2^) of the model to data from each state resulting from type A model fits. **(C)** Same as (A) but for type B model fits. **(D)** Difference in the goodness-of-fit (ΔR^2^) between type A and B fits. Negative values accuracy was greater in type B that type A fits.

Priors on model parameters dictating intrinsic dynamics (e.g., time constants, firing rate properties, etc.) were chosen using a combination of sources: (1) we preferentially used the Allen Brain Atlas data portal (https://celltypes.brain-map.org/) and retrieved properties derived from human cortical cells; (2) when parameters were not available in Allen Brain Atlas, we used the NeuroElectro database (https://neuroelectro.org/) as an alternative. For both databases, multiple estimates were available per parameter, and so we used the estimated mean and standard deviation to specify the respective expectations and precisions on (Gaussian) prior densities. Interlaminar connectivity was parameterized to match the same ratios of synaptic gains described in Bhatt et al. (2016). Prior covariances between parameters were assumed to be zero.

Systems of stochastic-delay differential equations (see supplementary methods IV) were solved numerically using a Euler-Maruyama integration scheme. For details of incorporation of finite transmission delays, and integration of the resulting system of stochastic-delay differential equations, see supplementary methods IV. We used an implementation of the sequential Monte-Carlo Approximate Bayesian Computation algorithm (SMC-ABC; Toni et al. 2009; West et al. 2021) to fit models. We take forward the maximum a posteriori (MAP) estimate (the mode of the marginal posterior distribution) of each parameter for additional simulations.

Model fits were assessed by the data used to fit them: *type A* – using the power spectra only; and *type B* – using both spectra and burst features (features described in section 2.4 “Data Features: Spectra and Distributions of Burst Amplitude/Duration”). We fit models to the group averaged data features and corrected the spectra to isolate peaks using a non-overlapping sum of Cauchy functions (see supplementary figure 3A and supplementary methods II). When fitting models across different motor states, the interstimulus interval (ISI) state was treated as a baseline, from which all other states were modulated. Thus, the ISI state was fit first using all free parameters (i.e., time constants, synaptic gains, sigmoid characteristics, properties of intrinsic and observation noise). The posteriors of the ISI state provided empirical priors for the remaining motor state models. These states were fit using a restricted set of free parameters incorporating laminar specific time constants, synaptic gains, sigmoid characteristics, and the slope/gain of 1/f^α^ innovation noise. All models were fit to the group averaged data features for each state.

### 2.7 Finding Parameters Responsible for Shaping Bursts

The posterior parameter estimates—under models of the motor cortex—were examined to identify parameters responsible for shaping burst properties. To do this, we individually manipulated the synaptic gain and gain parameters for the laminar specific inputs (a total of 18 parameters) on a logarithmic scale from −3 to +3 (equivalent to approximately decreasing or increasing the strength 20 times) in 24 steps. Each model was simulated for 48 seconds, and the following properties were estimated: the peak frequency of the spectrum, percentage change in power (from base model), mean burst amplitude, mean burst duration. Parameters correlating with each feature were then identified by estimating the Spearman’s rank correlation coefficient with the average of each feature (i.e., the expected value of the kernel approximation to the probability density function). This constitutes a sensitivity or contribution analysis: in other words, it assesses the degree to which changing synaptic parameters generate discernible differences in the space of data features.

As features may not correlate across the whole connectivity range due to, for example, the existence of bifurcations in the model, we computed correlations within a restricted range. The restricted range was identified by computing the Spearman’s coefficient between the parameter and mean feature value across all possible ranges, with a minimum window of 1/2 of the whole range examined (i.e., 12 steps in connectivity strength). Correlations were thresholded using a Benjamini-Hochberg correction to set the False Discovery Rate to 10%, and the range yielding the largest coefficient was selected. The correlation between average burst duration and parameter scaling was used to choose the range, as this feature was found to have the largest association with interlaminar connectivity. Correlations with the other three signal features (peak frequency, mean burst amplitude and interval) were taken within this parameter range. Finally, candidate parameters were found by examining the correlation coefficients. To identify parameters engendering changes in burst properties—but showing minimal effects on spectra—we looked for those exhibiting clear correlations with burst features but not with spectral frequency/power.

To link model parameters more concretely to dynamics, we constructed bifurcation diagrams from deterministic variants of the models using posterior (empirical) parameter estimates. Equilibria were identified from unique points in the steady state solution at which the approximate derivative was equal to zero. Stability of the equilibria was assessed by computing eigenvalues of the (delayed) Jacobian at each fixed point (David et al. 2006). For details, please see supplementary methods V.

### 2.8 Assessment of the Cortical Input/Output Fidelity and Relationship to Expression of Beta Bursts

Finally, we used the model to understand how parameters responsible for modifying stochastic burst activity may regulate a trade-off between beta modulation under spontaneous cortical activity versus that in response to exogenous input (e.g., as arising from sensory evoked potentials). To do this we delivered a train of inputs (modulations of asynchronous firing rate) to the middle pyramidal layer- the main recipient of thalamocortical afferents. We then assessed how this modulated beta bursts in deep cell layers – the predominant output layer of cortex (illustrated in figure 7). Inputs were given as a step function with bouts of length in seconds drawn randomly from a normal distribution with mean 500 ms and 150 ms standard deviation, and breaks drawn with mean 700 ms and 150 ms standard deviation. Inputs were multipliers on the stochastic firing rate and were set to 1x on the breaks and 3x (to test response to increase input rate) during bouts of upregulation. Fidelity of modulation was assessed by computing the Spearman’s correlation between the input (square wave of firing rate modulations) and output (square wave reflecting beta burst detection). We thus used this measure of input/output (I/O) fidelity to assess to what extent parameters known to regulate beta bursts also comodulate cortical transmission.

## 3 Results

### 3.1 Beta Burst Features in Motor Cortex are Modulated during Movement and are Better Predictors of Motor State than that of Spectra Features

Data features summarising the spectra (e.g., peak frequency, power in band), and characteristics of bursting activity (e.g., median burst duration/amplitude) were constructed from ECoG signals taken from the three datasets (see methods) and epoched to yield segments reflecting different motor states: rest/interstimulus intervals (ISI; colour coded in blue throughout), movement preparation (red), movement execution (green), and movement imagery (orange). Data were selected from a sensor close to motor cortex that exhibited the largest movement related beta desynchronization (see methods for selection criteria). Example time series from the different motor states are shown in figure 2A which show clear bursts of 14-30 Hz beta activity in data from the different states. Spectra in figure 2B demonstrate a clear movement related beta desynchronization in the group averaged spectra that is reflected in the change in 14-30 Hz narrow-band SNR from +18 dB to +11 dB from preparation to execution of movement (figure 2E; post-hoc t-test (40), P = 0.007). Changes were found in the wide band SNR (i.e., level of background noise indicating the overall signal quality) and corresponded to worsened recording quality during movement epochs (supplementary figure 1B). Beta desynchronization associated with movement is reflected also in a reduction in burst amplitudes (figure 2C and F; one-way ANOVA P = 0.011) and a shortening of beta burst durations (figure 2D and G; one-way ANOVA P = 0.004), although no significant changes were found in terms of the peak beta frequency or inter-burst intervals (supplementary figure 1 C and D, respectively).

To compare the predictive value of either spectral or burst features, we trained an ensemble of binary SVM classifiers to predict different motor states (supplementary figure 2). Decision boundaries (indicating > 50% prediction success) between all four motor states were present for classification with burst features, and AUCs of the receiver operating characteristics (ROCs) showed good predictive value (AUC > 0.80). In contrast, classifiers using only features derived from the power spectra could only separate features from movement preparation and movement execution states with AUCs > 0.5 (greater than chance level) and could not classify features derived from movement preparation or imagery states. These results suggest that, when using band restricted information (i.e., within 14-30 Hz), the properties of bursting activity can significantly augment the prediction of motor states from brain activity.

### 3.2 Burst Features are not Predicted by Linear Models of the Data

To further determine whether beta burst features reflect meaningful information about the underlying motor state, beyond that contained in the spectra, we compared empirical features with those computed from spectrally matched IAAFT surrogates (see methods). In figure 3, we show a comparison between empirical data features and the average feature derived from surrogate data (n = 25) for each of the motor states. By design, the surrogates matched well to the power spectra of the data (figure 3B and E). Differences between the distributions of burst amplitudes and durations computed from the data or from linear surrogates (figure 3C/F and D/G, respectively) show that both features deviate significantly (median R^2^ < 0.80) from that expected under linear assumptions. Comparisons of the goodness of fits (R^2^) to linear surrogates showed that deviations of burst duration distributions from linearity were not equal for each motor state (figure 3G, one-way ANOVA P = 0.001), with movement preparation and ISI states showing reduced R^2^ values when compared to movement execution. Similarly, burst durations exhibited significant changes between states (figure 3G, one-way ANOVA P = 0.002) with data from the ISI and movement preparation providing the greatest evidence for nonlinearity among all the motor states. These data suggest that burst features represent underlying nonlinearities in the data that are not captured in the power spectra alone. Further, states associated with ISI and movement preparation are associated with a higher degree of nonlinearity, especially when compared to movement execution. We next use a neural mass model to investigate the potential biophysical explanations for these differences.

### 3.3 Biophysical Models of Motor Cortex Fit Constrained to Fit Power Spectra do not Predict Distributions of Burst Features

We used the Sequential Monte Carlo Approximate Bayesian Computation (SMC-ABC) algorithm to fit a biophysical (neural mass) model of the motor cortex microcircuit to key data features (i.e., power spectra and distributions of burst duration/amplitude) from each of the four motor states. We fit the group averaged data features and further reduced spectra to their peaks using a sum of Cauchy functions (see supplementary figure 3D-F and supplementary methods II). To assess the value of the power spectra in predicting burst features, fitting procedures were split into two groups depending upon the data features used: *type A* - constrained exclusively using the spectra, or *type B* – constrained using a combination of the spectra and distributions of burst amplitude and duration (figure 4A). Samples of the simulated time series using posterior estimates, as well as the fitted features are shown in supplementary figure 3.

*Type A* models fit well to spectra (figure 4B; all states R^2^ > 0.95) but showed that spectra were not sufficient to predict burst features accurately. Further analysis of the fitted features (supplementary figure 3E and F) showed that predicted distributions of burst amplitudes were attributable to smaller amplitude bursts than those observed in the experimental data, and burst durations were shorter than predicted in the case of ISI and movement preparation (blue and red, respectively; R^2^ < 0.90). However, *type A* fits were sufficient to accurately recover the empirical distributions of burst amplitude in movement execution/imagery (figure 4C; green and orange, R^2^ > 0.90).

In contrast, *type B* fits demonstrate that the model parameters could reproduce burst features (figure 4C), with a median fit of ∼95% for all features. Complementary to the analyses of feature nonlinearity in figure 3, we show that the ISI and movement preparation (the motor states exhibiting the highest degree of nonlinearity) gained the most (in terms of accurate predictions) from the explicit inclusion of burst features (difference of *type A* and *B* fits shown in figure 4D). In contrast, for data from movement imagery and execution there was less gain in accuracy when explicitly incorporating burst features.

The inadequacy of *type A* fits in predicting burst features (withheld from model inversion) suggests that burst characteristics are the product of circuit mechanisms (and associated biophysical parameters) that are either independent or at least only weakly associated with those governing the power spectral amplitude and implies that features summarising temporal patterning of bursts are important for informing neural models. Furthermore, burst features from periods of ISIs and preparation appear most different from those predicted using *type A* fits out of all of the other motor states. In the next section we aim to identify parameters of the fitted microcircuit models of motor cortex underlying these changes in burst properties.

### 3.4 Analysis of Parameter Modulations Between Motor States and Correlations with Burst Features

Parameters of the fitted models exhibited significant deviation from the empirical priors provided by the model fit to movement preparation (i.e., the baseline state), indicate changes in cortical microcircuity between motor states predicted by the model (figure 5A and B). Parameter estimates based on ISI, movement imagery, and movement execution data showed significant changes in drive to inhibitory interneurons (II input gain), with the latter two showing an increase in inhibition. Movement preparation, execution, and movement imagery were also associated with changes in self inhibition of deep layers (DP → DP).

**Figure 5.**
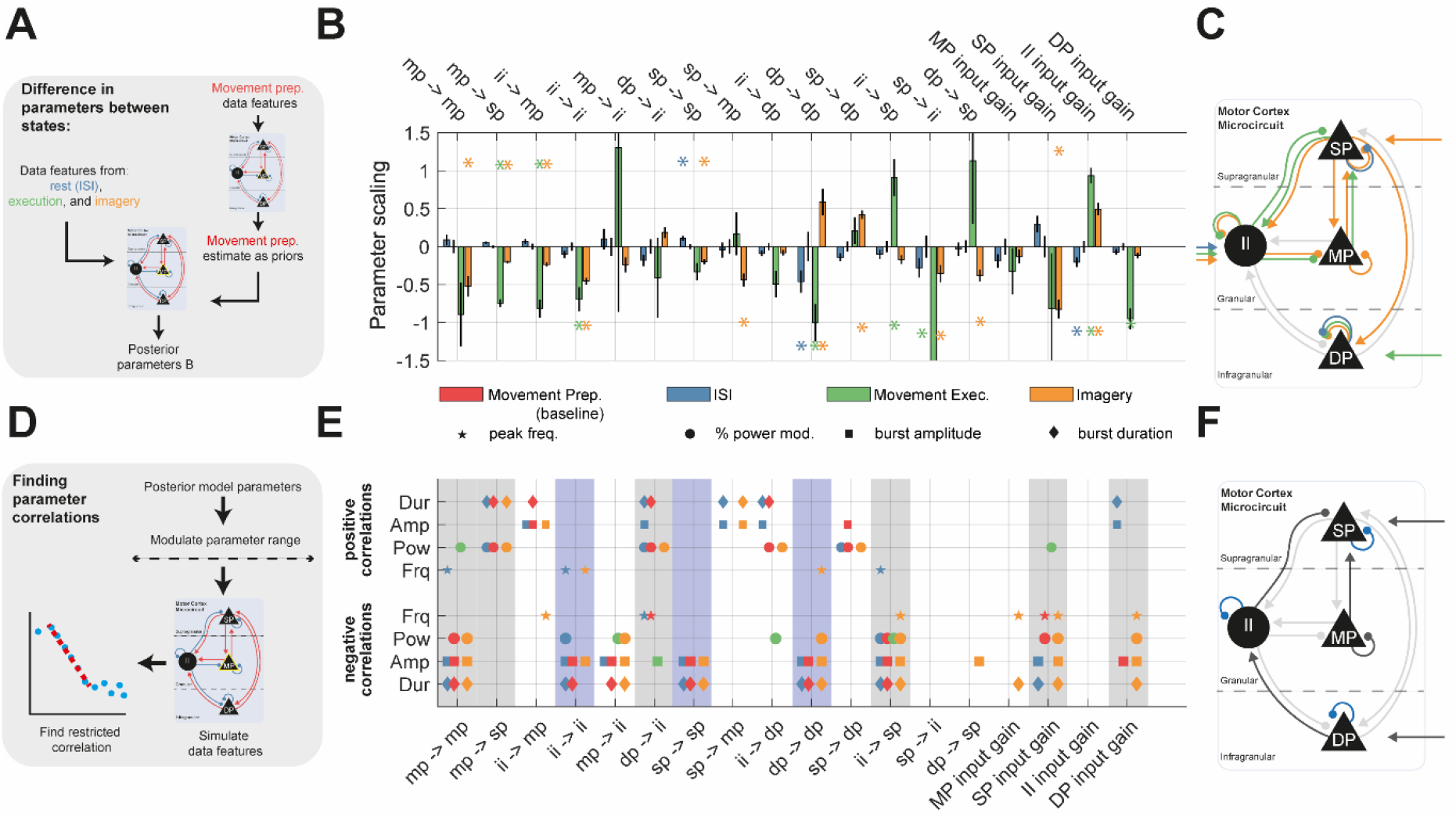
Results of the motor cortex model fits to ECoG data from motor tasks. Analysis shows posterior model estimates, as well as modulations in parameters from the baseline condition (movement preparation.), as well as correlation analysis of circuit parameters with the statistics of spectral and burst features resulting from posterior simulations. **(A)** Parameters of the model of motor cortex microcircuit were estimated from fits to group averaged data features from all four motor states using ABC-SMC. **(B)** Changes in parameters from the baseline, movement preparation state (red-zero for all parameters indicating usage of empirical priors for the remaining states) show statistically significant modulations (posterior Z-test P < 0.05, indicated by asterisk), particularly for features estimated from movement execution and imagery. **(C)** Connections exhibiting a significant modulation are shown on the colour coded circuit diagram. **(D)** Modulations in parameters were estimated by first fitting to movement preparation data as a baseline state (using a wider set of free parameters, see methods), and then using these as empirical priors on the remaining models (using a smaller set of free parameters, see methods). **(E)** Parameters of the posterior models dictating interlaminar connectivity, and laminar specific inputs were then systematically examined for correlation with different data features. Correlations were performed on a restricted range (see methods). Parameter significance was determined using False Discovery Rate correction (10%). Grey bands highlight parameters that modulated both power and burst features. Parameters in light grey reflect those predominantly acting on burst features. **(F)** Connections and inputs exhibiting a significant correlation with either spectral and burst features (highlighted in grey) or exclusively burst features (blue) are shown on the colour coded circuit.

To identify the parameters responsible for shaping beta burst features, we systematically altered interlaminar connection strengths and input gains, and then applied a restricted-window correlation analysis (see methods) to detect co-modulation of the parameter with the predicted spectral frequency, beta power, mean burst duration, or mean burst amplitude (figure 5D). The results in figure 5E show that common parameters affect these data features in models fitted across the motor states. Parameters modulating both burst and spectral features (highlighted in grey in figure 5E) included: MP self-inhibition; II → SP; SP input gain. With respect to beta burst features, three parameters were found to predominantly modulate burst amplitude and durations (highlighted in light blue in figure 5E). Interestingly, all three parameters correspond to self-inhibitory connections for SP, DP and II. To investigate how these parameters shape beta dynamics, we chose an example parameter—DP self-inhibition gain—that we took forward for further analysis. This was because: (A) it shows strong modulation between motor states (figure 5B); and (B) it negatively correlates with both burst amplitude and duration but exhibits only limited effects on spectral peak frequency or power (figure 5E).

### 3.5 Analysis of the Fitted Models Demonstrates that Bursting Intermittencies are Shaped by Dynamical Stability of the Motor Cortex

We used DP self-inhibitory gain as a control parameter to investigate its effects on temporal dynamics in the simulated model (figure 6A) and to construct bifurcation diagrams (figure 6B). To understand how this control parameter may change the system’s response to small perturbations (such as that provided by small amplitude noise), we performed a stability analysis of the estimated equilibria.

**Figure 6.**
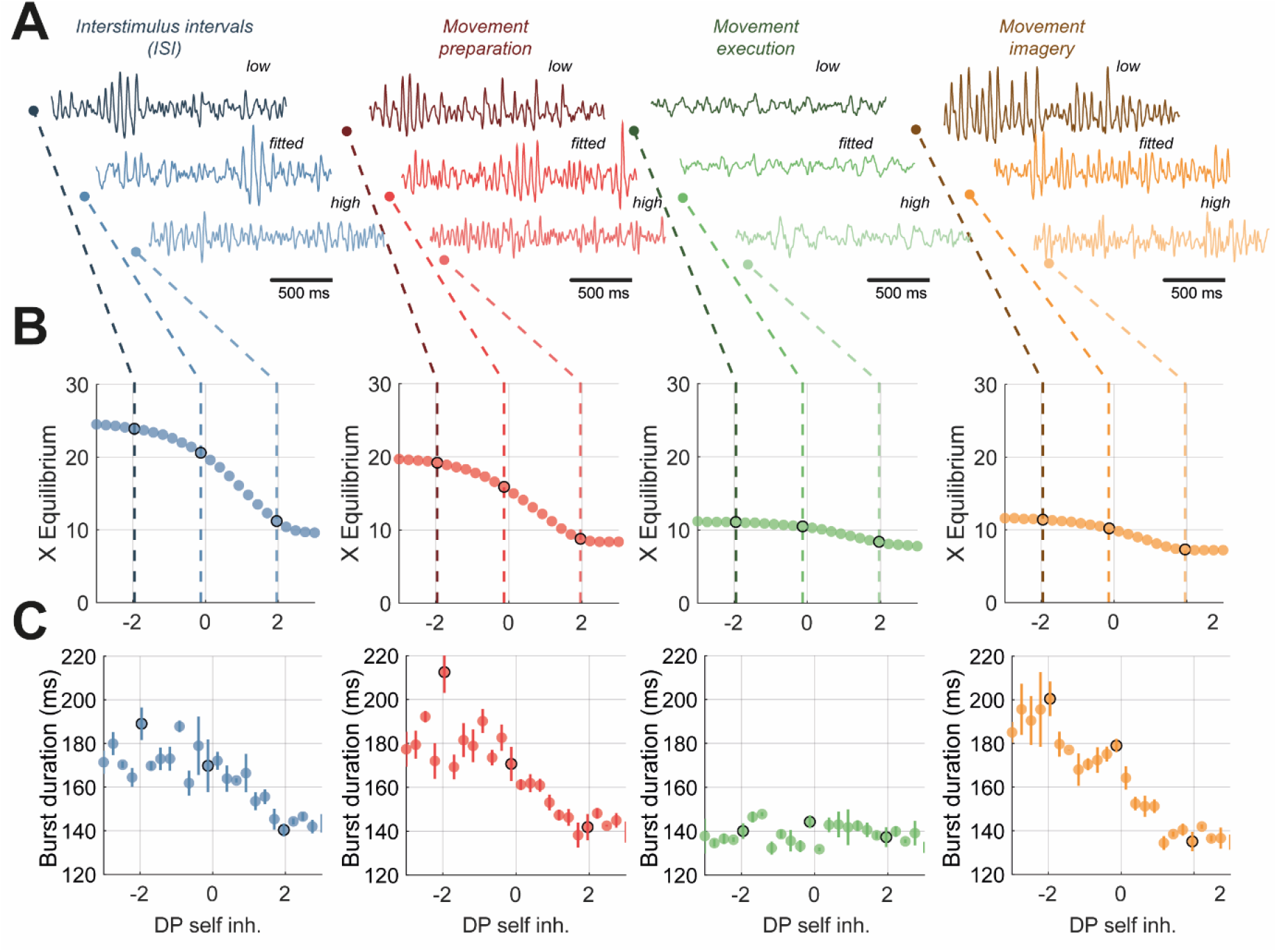
Detailed model analysis of bifurcation diagrams associated deep pyramidal layer (DP) self-inhibition strength and corresponding correlations with signal features in terms of burst duration and nonlinearity. The level of deep layer self-inhibition was taken forward as a control parameter following from the correlation analysis presented in figure 6F. Simulations were performed on a range of parameter values spanning −3 to +3 (log scaling from posterior). **(A)** 1.5 seconds of sample data simulated from each model of a motor state at either low (−2 scaling), fitted (0 scaling), or high (+2 scaling). **(B)** Bifurcation diagrams estimated from the deterministic variant of the model (see methods). Dashed lines indicate correspondence between stochastic model dynamics and level of control parameter. All states show fixed point dynamics. **(C)** The median burst duration is plot against the strength of DP cell input. All states excluding movement execution indicate existence of negative correlation between control parameter and burst duration. For bifurcation analysis and analysis of DP self-inhibition effects on signal nonlinearity (using IAAFT2 surrogates) please see supplementary figure 5.

**Figure 7.**
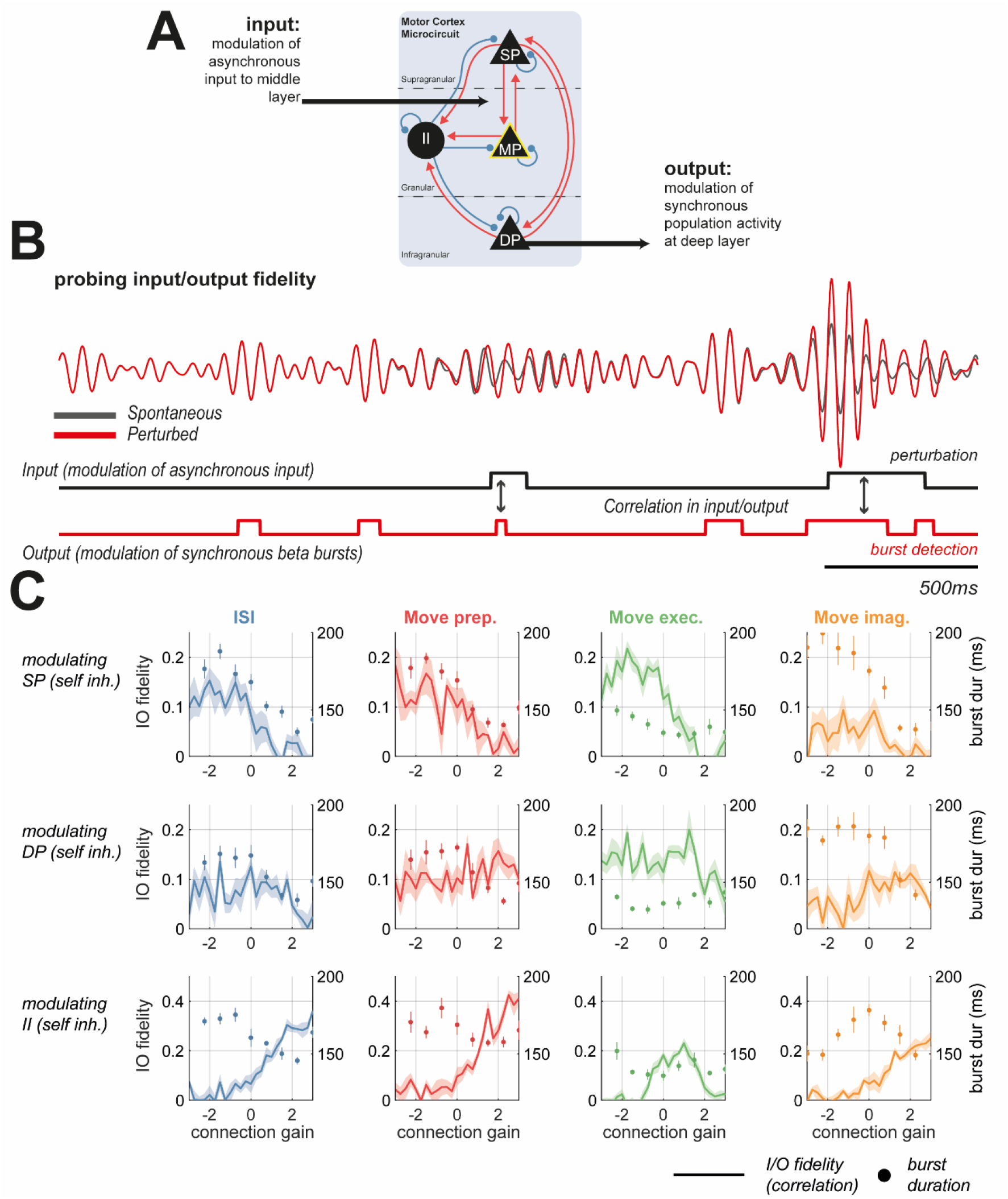
Parameters responsible for modulating burst properties do not uniformly alter the fidelity of synchronous cortical responses to exogenous inputs. **(A)** To probe the fidelity of cortical beta responses to changes in exogenous input, fitted models were used in an in-silico experiment. Asynchronous (stochastic) inputs to the middle layer were modulated with a square wave of random intervals. Beta burst detections in signals simulated in deep cell layers were taken as the outputs. The total “fidelity” of input/output (I/O) transmission was estimated using the rank correlation/mutual information between these two square waves. **(B)** Example waveforms of the spontaneous (unperturbed; grey) activity, overlaid with perturbed (in red) activity matching the perturbation (i.e., modulation in noise to middle layer) seen below (black square wave). The output of the system matches the beta burst detections (red square wave) **(C)** Self inhibition in superficial (SP), deep (DP), and inhibitory interneurons (II) layers negatively correlates with burst duration (given on right axes; dots; see figure 5 and 6). However, modulations in I/O fidelity (shaded lines) do not align with burst duration: SP self. decreases I/O fidelity, DP self. exhibits no correlation with I/O fidelity, and II self. increases fidelity.

All models—across the motor states—exhibited stable fixed-point dynamics for at least some of the range investigated. Sustained oscillatory activity is observed in models fit to the data recorded during ISI, movement preparation and movement imagery states. This oscillatory activity was present for DP self-inhibition in the range −2.5 to +2.5 (log-scaling factor; shown in traces figure 6A, in blue, red, and yellow). Analysis of the deterministic system showed that this change correlated with a reduction in amplitude of the system’s equilibria (figure 6B) and a change in the stability of the equilibria (analysis shown in supplementary figure 4B). Corresponding intermittencies in beta rhythms were graded, with burst durations shortening continuously as DP self-inhibition was increased (figure 6C; blue, red, and yellow). In the model fit to data recorded during movement execution (in green), there was no periodic behaviour in the simulated traces generated by the model (figure 6A, green) and DP self-inhibition showed no modulation in the equilibria or eigenvalues of the system (figure 6B and supplementary figure 4). There was also no modulation in burst duration with this parameter. We also analysed changes in feature nonlinearity (using comparison to IAAFT2 surrogate method introduced in section 3.2) but found that the high variance of the estimator impaired any detection of modulation by the control parameter (supplementary figure 4A). These analyses demonstrates that the duration of temporal intermittencies of beta rhythms in the model, can be explained by the effects of biophysical parameters on the system, in this instance, DP self-inhibition reduces the magnitude of the oscillatory response to perturbation, such as that occurring due to noise in the stochastic model.

### 3.6 Statistics of Beta Burst Expression Cannot Reliably Inform the Receptivity of the Cortex to Exogenous Inputs

Finally, we investigated the hypothesis that cortical beta burst properties reflect a trade-off between integration of spontaneous endogenous activity, versus that arising due to structured exogenous inputs (Karvat et al. 2021) (i.e., from sensory or higher order thalamus). In figure 7A and B we illustrate an in-silico experiment conducted on the models fit to different motor states in which we delivered patterned modulations of asynchronous (i.e., noisy) inputs to the middle layer of cortex (the main recipient of thalamic projections). We considered beta burst detections in deep layer (the main projection layer of cortex) as the cortical output. We then measured the correlation between the input and output as an estimate of transmission fidelity. This analysis was repeated separately for all three self-inhibition strengths known to negatively correlate with beta burst amplitude and duration (shown in figure 5): SP self., DP self., and II self.

The results in figure 7C show that whilst strengthening all three of these parameters decreased mean burst duration (right axes; shown by dots), the relationship with I/O fidelity was not consistent between different self-inhibitions. For instance, in the models fit to ISI data, SP self-inhibition associated shortening of bursts correlated with a decrease in I/O fidelity. The opposite was true for modulations of II self-inhibition gain. These data suggest that burst statistics are not sufficient to infer the integration of endogenous and exogenous information in the cortex as shortening of bursts can be associated with both increased and decreased translation between exogenous inputs and modulation of beta frequency transients.

## Discussion

### 3.7 Summary of Findings

Temporal dynamics of rhythmic activity in the brain contain significant information regarding cortical information processing. Here, we have shown that motor states can be decoded from electrocorticography using features computed from narrow-band beta activity (figure 2). Our results show that these features aid classification (supplementary figure 2) and arise from signal nonlinearities that are not detectable in the power spectrum (figure 3). Further, evidence for nonlinearity was found to be greatest in data recorded during rest and movement preparation, indicating that the increase in information, beyond that available in the spectrum, and contained in the distributions of burst amplitude/duration, is highest in these states. Using a neural mass model, we then delved into the potential mechanisms and their functional significance. As expected, we found that neural mass models fit exclusively to spectra were not sufficient to accurately recapitulate the features of cortical beta bursts (figure 4). Analysis of the fitted model parameters between motor states found that burst properties could be modulated by specific interlaminar couplings, and independently of spectral amplitude or frequency (figure 5). These parameters were predominantly self-inhibitory connections to deep, superficial, and inhibitory interneuron populations. Using deep self-inhibition as an exemplar control parameter, we showed how changes to the equilibria and dynamical stability of the deterministic model, could in turn shape the properties of spontaneous beta bursts when noise was added (figure 6). Finally, using simulations of the fitted models, we showed that changes in burst duration and amplitude cannot reliably infer receptivity of the cortex to input, as the relationship was dependent upon the specific connection responsible for altering bursting (figure 7).

### 3.8 Intermittencies in Bursts can Discriminate Brain States Associated with Movement

Transient fluctuations in neural oscillations can contribute to the understanding of the organization of brain activity (Bonaiuto et al. 2021; van Ede et al. 2018; Feingold et al. 2015; Lundqvist et al. 2016; Sherman et al. 2016; Shin et al. 2017). Transients in beta oscillations, the focus of this study, are found in healthy sensorimotor cortex (Feingold et al. 2015; Hannah et al. 2020; Little et al. 2019; Rule et al. 2017; Wessel 2020), and also play a prominent role in Parkinsonian electrophysiology (Cagnan et al. 2019; Tinkhauser et al. 2017b). Quantification of these intermittencies is beginning to build a taxonomy of bursts by identifying changes associated with different brain states and diseases (Deffains et al. 2018; Enz et al. 2021; Khawaldeh et al. 2020; Shin et al. 2017; Torrecillos et al. 2018). The discrimination of brain states by temporal features, as well as their transitory nature, makes them attractive targets for closed-loop approaches to neuromodulation, for instance using either beta frequency (Little et al. 2016; Tinkhauser et al. 2017a), or theta and gamma (Kanta et al. 2019; Knudsen and Wallis 2020) biomarkers.

The results reported here support this approach, by providing direct evidence that quantification of burst duration and amplitude, from narrow-band information can aid classification of motor states, in a way that is superior to that achieved when using spectral measures of beta power or peak frequency alone. Notably, we were able to discriminate between periods of rest and movement preparation, despite similar beta SNR observed across these states. These burst features are good candidates for control signals in closed loop neuromodulation, as they can be readily computed from narrowband data such as that available on current sensing devices such as Percept (Van Rheede et al. 2022) and they are known to be modulated by deep brain stimulation (Pauls et al. 2022). Additionally, motor state discrimination was enhanced compared to linear surrogates, with the degree of nonlinearity largest during rest and movement preparation (figure 3). This technique has previously been deployed to show that Parkinsonian beta bursts are more nonlinear when compared to a medicated control state (Duchet et al. 2021). This suggests the possibility that biomarkers relating to signal nonlinearity can also form the basis for novel closed loop control algorithms (Jelfs et al. 2010) for neuromodulation.

### 3.9 Mechanisms and Functional Implications of Bursts in the Motor Cortex

If the statistics of bursts in rhythmic neural activity are discriminating features of brain states, then they may provide a window into the underlying changes in the generative neural circuitry. Existing models show that interactions between synchronous subthreshold inputs to proximal and distal dendrites of pyramidal neurons can explain high amplitude, short duration bursts of beta recorded in sensorimotor cortex (Bonaiuto et al. 2021; Sherman et al. 2016). Strong inputs to distal dendrites may then halt information processing by recruitment of inhibitory interneurons in the supragranular layers (Jones et al. 2009), that can lead to a reduction in pyramidal firing rates following cortical beta bursts (Karvat et al. 2021). Our model also suggests that the strength of projections from superficial to deep lamina is an important determinant of total beta power, yet this parameter does not explain changes in the temporal dynamics of bursts. It is likely that the high amplitude waveforms chosen in these previous studies to maximize signal SNR, form only a subset of the total beta activity as there is good evidence for motor cortical bursts lasting > 300ms in duration (Seedat et al. 2020). Thus, a focus on high amplitude beta events may occlude alternative mechanisms by which recurrent and delayed interlaminar interactions may either seed the genesis of beta bursts and/or sustain them across multiple cycles. For instance, our work suggests an important role for laminar specific inhibitory interneuron activity, with deep layer self-inhibitory loops acting to curtail burst durations.

As changes in temporal patterning of beta activity between motor states are ascribable to alterations in interlaminar connectivity, it thus follows that the amplitude modulation of beta oscillations may reflect changes in the response to driving inputs to the cortex. The cortex is known to exhibit context dependent changes in interlaminar propagation and laminar specific inputs (Kirchgessner et al. 2020; Takeuchi et al. 2011) yet limited information is known regarding the changes occurring during movement (Inagaki et al. 2022), and even less about how this relates to the frequency of activity. Our simulations demonstrate that input/output relationships between exogenous modulations in firing rates and beta entrainment may change between brain states. However, there was no consistent finding that burst properties (i.e., burst elongation) corresponded changes in integration of exogenous inputs (figure 7), as the relationship changed dependent upon whether bursts were elongated by superficial or inhibitory inhibition, for instance. Thus this model is unable to provide evidence in support of the idea that spontaneous beta bursts in sensorimotor cortex reflect a competition with sensory evoked potentials (Karvat et al. 2021).

In the cases that beta bursts do reflect sensory gating (Van Ede et al. 2011; Limanowski et al. 2020; Spitzer and Haegens 2017), then high amplitude or elongated beta events arising from increased stability of beta generators (as suggested by our analysis in figure 6) could reflect a down weighting of sensory inputs in favour of maintenance of the existing motor program and enhanced robustness to sensorimotor “noise” (Cocchi et al. 2017). Our simulated experiment (presented in figure 7) suggests that the fidelity of cortical responses to external perturbation should change dependent upon motor states. This could be validated, for instance, by providing patterned optogenetic stimulation to specific layers, and then measuring the fidelity of the cortical response.

### 3.10 Model Inference and Intermittent Dynamics

This work also provides evidence that power spectra alone may contain insufficient information to accurately constrain parameters of nonlinear and/or stochastic models. Existing dynamic causal models of large scale temporal dynamics such as Parkinsonian beta bursts (Reis et al. 2019) or epileptic seizures (Rosch et al. 2018) appeal to fast-slow separation of time scales (i.e., the adiabatic approximation) in which changes in dynamics (i.e., bursting to quiescence) can be approximated by a model of fast (i.e., oscillatory) dynamics, with slow variables regulating the transition between states (Jafarian et al. 2021). In a similar vein, many phenomenological or statistical models describe bursts as a transition between discrete dynamical states (Heideman et al. 2020; Seedat et al. 2020). Other modelling approaches, such as that of Sherman et al. (2016), described above, take well constrained compartmental models that can describe high amplitude beta events, albeit with a specific pattern of input.

In this paper we take a different approach and treat bursts as the product of stochastic “quasi-cycles” that arise from noise driving a stable system such as a damped oscillator (Powanwe and Longtin 2019), that exhibit amplitude envelopes that can be modelled in terms of a drift-diffusion process (Duchet et al. 2021). Thus we use a model incorporating the full nonlinear transfer functions, and fit parameters of the resultant stochastic differential equations (West et al. 2021). Given the full breadth of information summarised by both the spectra and distributions of burst features, these models can well describe temporal dynamics of ECoG data in a parsimonious way without needing to appeal to modelling multiple states separately.

The distinction between generative models in which synaptic parameters fluctuate slowly and our model based upon stochastic dynamics speaks to an important distinction between explanations for itinerant dynamics of which beta bursts provide a good example. Technically, the first kind of generative model rests upon *structural instability*, where the itinerant changes in fast neuronal dynamics—and ensuing transients—are generated by changes in the fixed points of a system with the parameters of the equations of motion. In contrast, the second kind of generative model relies upon *dynamical instability*; namely, unstable (or weakly stable) fixed points to produce transient dynamics. This formal distinction has importance for understanding the biophysical mechanisms that generate bursts in population activity, as well informing stimulation approaches that aim to modulate them. For instance, in the case that bursts are the direct product of slow changes in neural circuits (i.e., invoking neural plasticity), then stimulation should directly target these mechanisms, whereas in terms of dynamical instability, stimulation can be patterned to with the aim of suppressing transient burst activity, or disrupting neural states that preclude them. Formally, this question could be answered in terms of a Bayesian model comparison between generative models incorporating either dynamic and structural instability.

### 3.11 Limitations

A major problem when investigating changes in temporal dynamics between brain states arises from potential confounds that arise from the trivial effects of changes in signal to noise. We note that we found changes in the wide-band SNR (i.e., the overall signal quality - compared to the amplifier noise floor) between states (supplementary figure 1). However, the variance of the wide-band SNR between subjects was very high and showed smaller effect sizes than that observed when comparing distributions of burst amplitude and duration, suggesting that SNR was not the main contributing factor. Further, alterations in burst amplitude did not correlate with either wide- or narrow-band SNR. The segregation in burst amplitude and duration effects between states was also sufficient to provide superior classification of states to that achieved when using SNR. Further, bursts were defined using a window-specific threshold, which prevents burst properties from predominantly reflecting SNR differences-a problem that is encountered when using a common (i.e., across states) threshold (Schmidt et al. 2020). The robustness of using a fixed threshold of 75^th^ percentile is well supported following reports that specific threshold values do not qualitatively change outcomes of burst analyses (Lofredi et al. 2019; Tinkhauser et al. 2017b).

To ensure good data quality, we applied stringent selection criteria (described in methods section 2.2) that lead to the rejection of significant portions of the available data. Data quality vs beta desync. Get both. Focus on robust effects

The existence of non-identifiability in models (i.e., a redundancy in parameter to output mappings) will always limit the degree of confidence with which parameter estimates can be interpreted. In terms of Bayesian models such as that presented here, the existence of prior densities over parameters can reduce these concerns to some extent, by providing an *a priori* restriction on the values to which parameters may take. This comes with the caveat that the mechanistic conclusions must only be interpreted in terms of the model architecture (the product of a previous model comparison study in (Bhatt et al. 2016)) and the specified priors (many of which are ascertainable from electrophysiological studies: see supplementary table).

Lastly, model inversion with Approximate Bayesian computation is susceptible to issues arising due to insufficiency of the summary statistics (i.e., the power spectrum, or distributions of burst duration/amplitude used here). More complete descriptions may be achievable with the bispectra (i.e., the Fourier transform of the third-order cumulant) (Halliday et al. 1995). Although there are dynamic causal models of cross-frequency coupling—implicit in the nonlinear mechanisms that underwrite dynamical itinerancy (Chen et al. 2009; Friston et al. 2006)—they are not generative models of bispectra, or indeed the statistics of bursts or transients. The results of the current study clearly call for development of generative models of these kinds of data features.

## 3.12 Conclusions

This work provides significant evidence that the temporal properties of bursting intermittencies in brain rhythms contain unique information about the underlying circuits that generate them, beyond that more conventionally inferred from the power spectra of electrophysiological data. Furthermore, we have shown that burst features are nonlinear and are not simple predictions of the power spectra. Using a model of motor cortex microcircuitry, we show that bursts can arise from stochastic dynamics, with properties that are predominantly modulated by local laminar specific inhibitory loops. We have shown that this has important consequences for understanding information processing in cortical microcircuits, although simulations exhibit a non-trivial relationship between burst duration and amplitude versus the responsivity of the cortex to exogenous inputs. These findings inform novel paradigms to understand the role of external perturbations such as electrical brain stimulation, in manipulating cortical computations when in the presence of spontaneous fluctuations in neural rhythms.

## 4 Acknowledgements

SFF Acknowledges personal funding support from the UCLH Biomedical Research Centre (BRC).

## 6 Supplementary Figures

### 6.1 Supplementary Figure 1

**Supplementary Figure 1.**
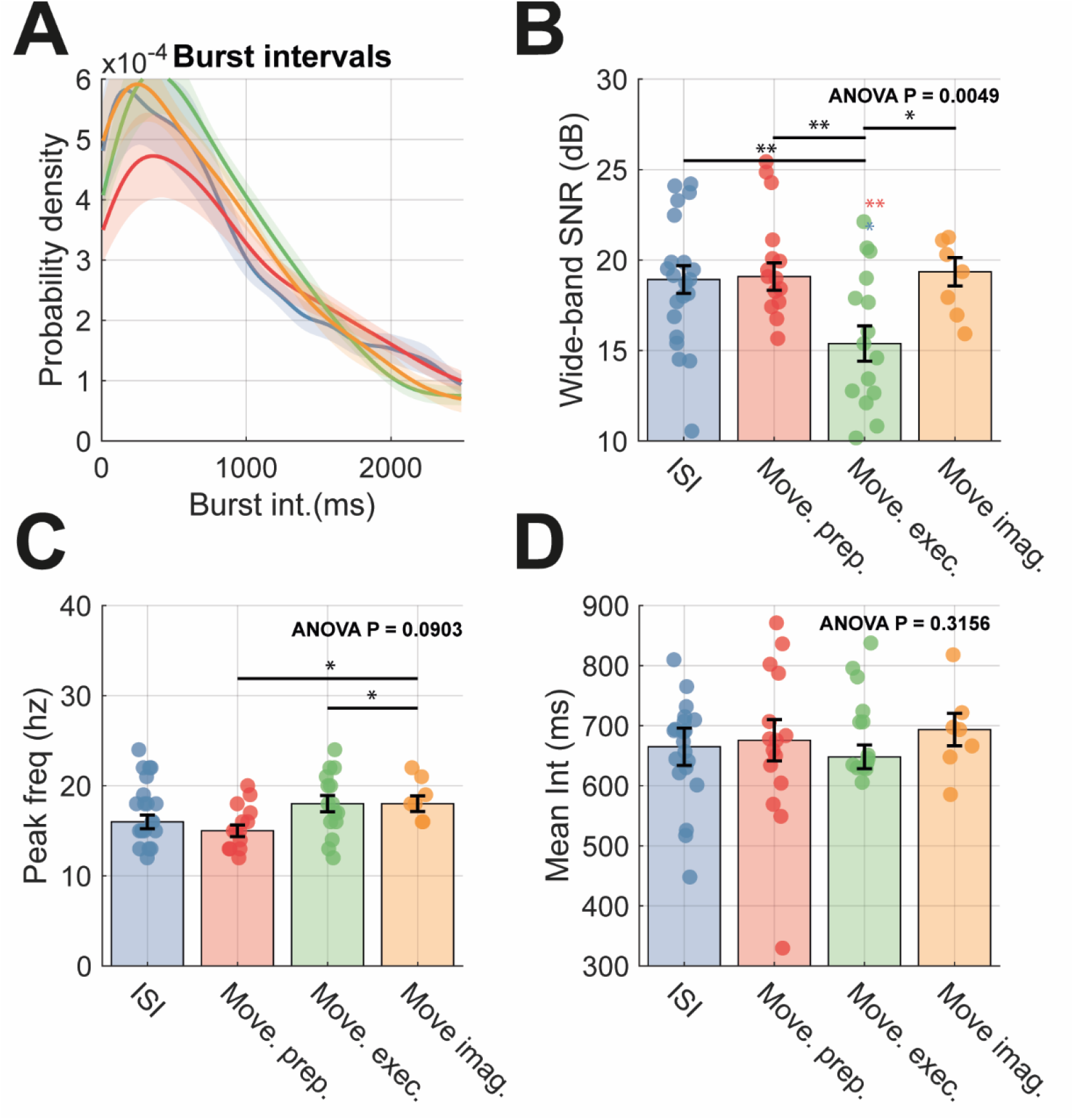
Additional ECoG signal features compared between motor states. **(A)** Probability densities of interburst intervals. **(B)** Bar chart to compare changes in the wide-band SNR of the selected ECoG channel. **(C)** Same as (B) but for peak beta frequency. **(D)** Same as (B) but for the mean interburst intervals.

### 6.2 Supplementary Figure 2

**Supplementary Figure 2.**
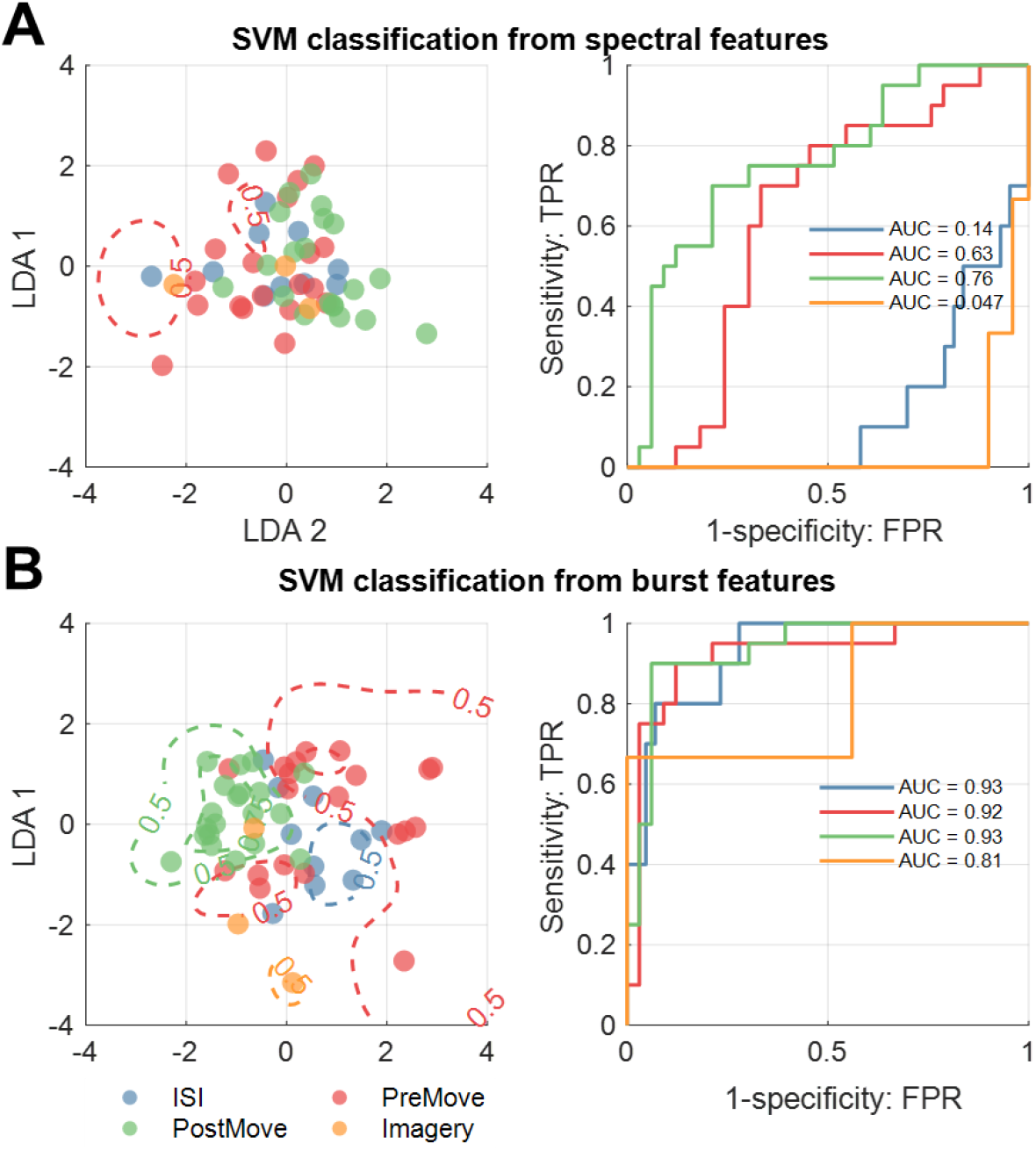
Classification of movement states is superior when using beta burst features over that performed when using spectral features only. **(A)** (left) Features of the ECoG power spectra (n=3) were projected onto a two-dimensional space using linear-discriminant analysis (LDA). Classification was then performed using ensembles of support vector machines on the first and second components of the LDA. The classification boundaries for each state are overlaid on the scatter plots of LDA features, at P = 0.5; and P = 0.75. (right) The receiver operating characteristics of each binary classifier are shown, with the area under the curve is inset. **(B)** Same as for (A) but when using burst features (n=6).

### 6.3 Supplementary Figure 3

**Supplementary Figure 3.**
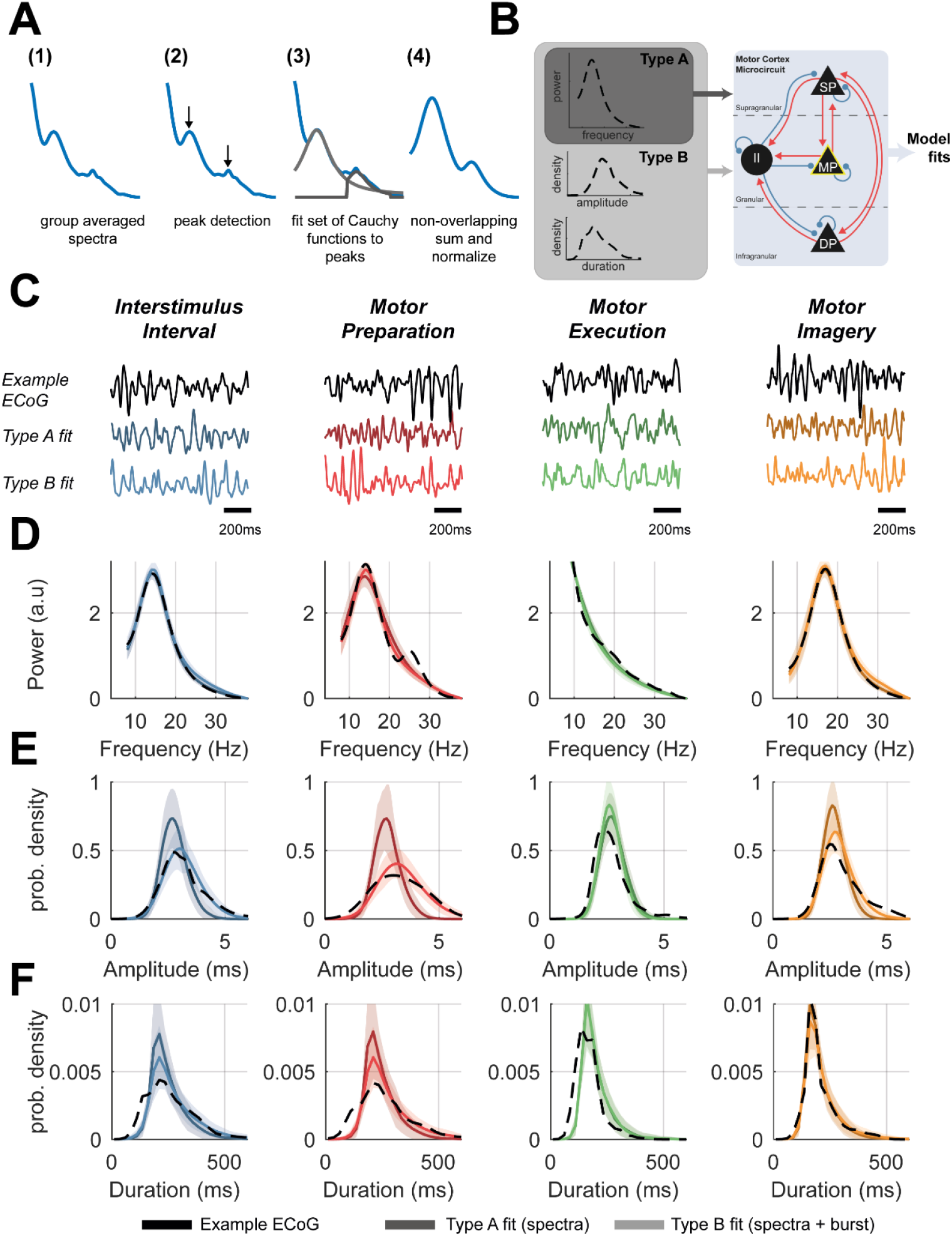
Summary of model fits of motor microcircuit model to group averaged data features across motor states. **(A)** Illustration of spectral preprocessing performed to isolate main peaks of spectra from 1/f background. **(B)** Schematic of the motor cortex microcircuit model. Each black node represents a neural mass that is coupled with either excitatory (red) or inhibitory connections (blue). There are three pyramidal cell layers: superficial (SP), middle (MP), and deep (DP), plus an inhibitory interneuron (II) population. Model parameters were constrained using either pre-processed spectra (type A) or both spectra and burst features (type B). **(C)** 1.5 second of example empirical data is shown from each motor state (top; dark shade), alongside those simulated from the posterior type A (middle; medium shade), or type B (bottom; light shade) fits. Data is shown from the interstimulus interval (blue), movement preparation (red), movement execution (green), and movement imagery (orange). Data features from the posterior model fits are shown for: **(D)** power spectra, **(E)** distributions of burst amplitudes, and **(F)** distributions of burst durations.

### 6.4 Supplementary Figure 4

**Supplementary Figure 4.**
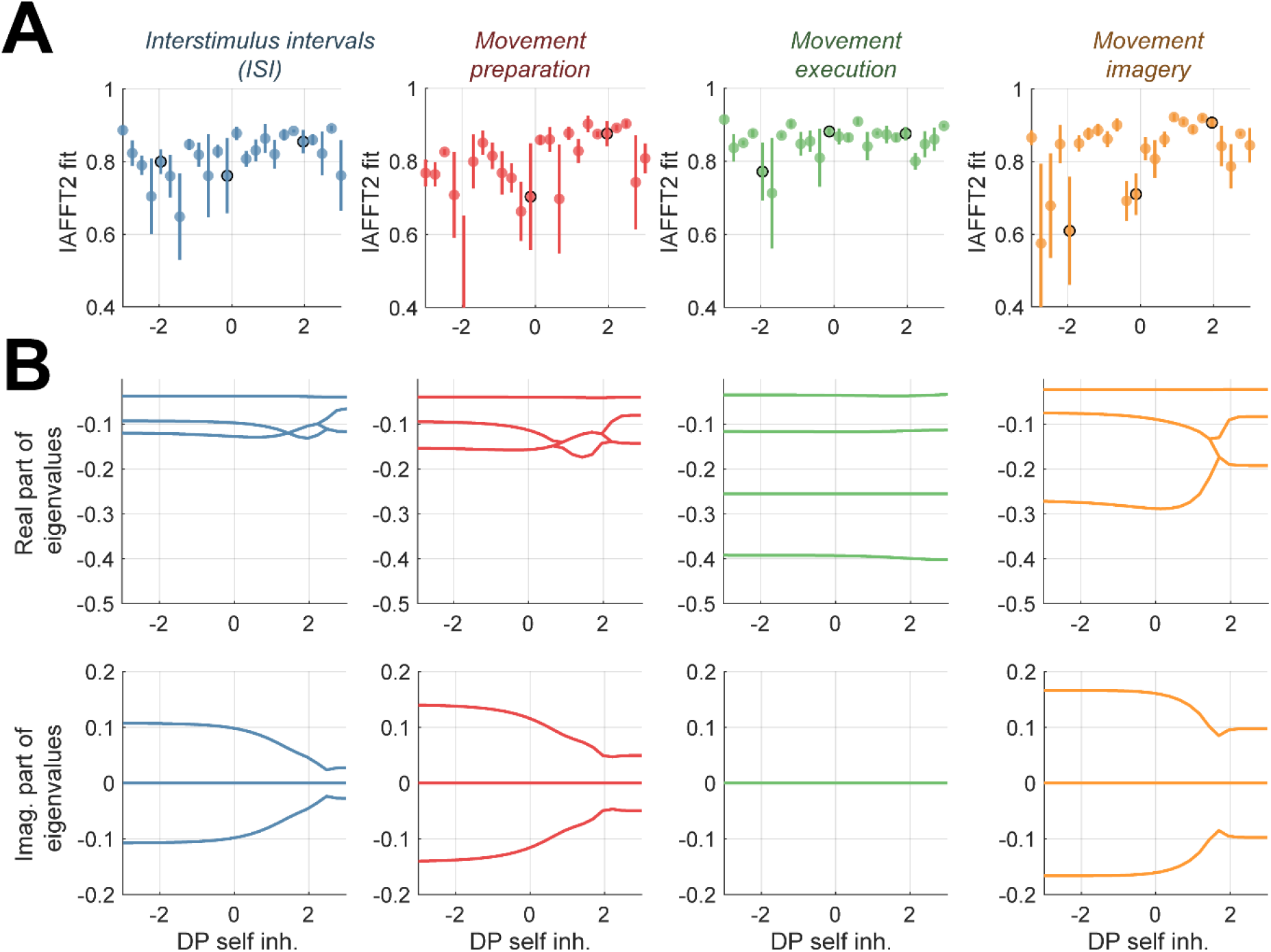
Bifurcation diagrams of system shown in figure 6. **(A)** The goodness of fit between burst duration distributions estimated from simulated data and linear surrogates indicates that the degree of nonlinearity in the signals is anticorrelated to changes in the burst durations. **(B)** Birfurcation diagrams indicate changes in either the real (top row) or imaginary (bottom row) components of the eigenvalues computed from the delay corrected Jacobian for each of the equilibria in each of the four models (parameterised to fit data from each of the four motor states).

### 6.5 Supplementary Methods

#### 6.5.1 Supplementary Methods I –Wide/Narrow-band SNR Calculations

#### 6.5.2 Supplementary Methods II – Spectral Reduction

Spectra were preprocessed prior to ABC model fitting in order to remove the aperiodic 1/f background such that fits were focussed on beta band activity. Peaks in the power spectrum in the beta frequency range were found using the *findpeaks* algorithm implemented in MATLAB. Prior to peak finding, spectra were smoothed with a 5 Hz wide Gaussian kernel. Inflection points (i.e., the troughs separating peaks) were then determined by finding the nearest sign change of the approximate derivative (difference) from each peak. This then defined the frequency range over which a Cauchy function was fit. This procedure was formed for each peak. The composite spectra were then formed from the non-overlapping sum of each fitted model.

#### 6.5.3 Supplementary Methods III – Definition of Bursts

Bursts were defined by setting a threshold on the bandlimited envelope. The filter passband was set at ±5 Hz of the peak frequency and implemented using a zero-phase FIR filter. Filtered data were then Z-normalised. The analytic signal was constructed using the Hilbert transform to estimate instantaneous amplitude. Bursts were defined as periods exceeding the 75^th^ percentile of this envelope and the minimum burst length was set to 2 periods of a 30 Hz oscillation (the upper limit of the band). Bursts found at the boundaries of epochs were discarded from the analysis. Burst amplitudes were taken as the maximum of the envelope within each burst, whilst burst duration reflects the amount of time that the envelope exceeds the threshold. Inter-burst intervals represent the time spent sub-threshold between each event. To summarise burst features, we estimated distributions of burst duration, amplitude, and inter-burst intervals using binned histograms. Distributions were then estimated using a kernel density estimate of the probability density function specifying a standard normal function for the kernel.

#### 6.5.4 Supplementary Methods IV –Model Formulation

The model uses the firing rate equations (Vogels et al. 2005; Wilson and Cowan 1972) constructed with the same architecture outlined in (Bhatt et al. 2016). The average firing rate of each laminar population (middle *MP*, superficial *SP*, inhibitory interneuron *II*, deep *DP*) is given by the following state equations:

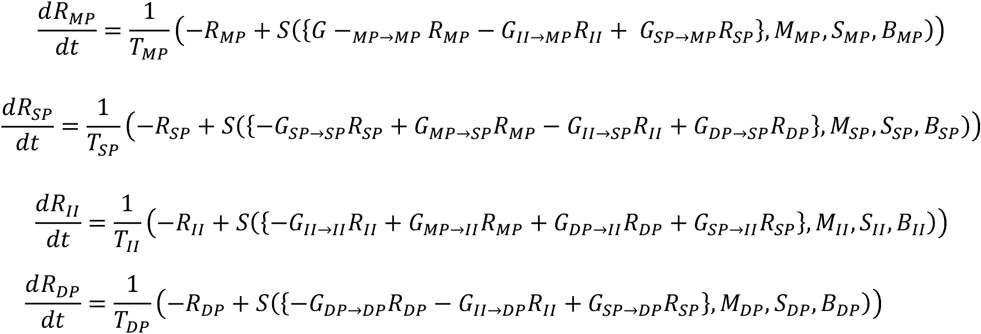

Where *T* gives the population time constant, *G* gives the weight of the (delayed) synaptic connection, and *S(I,M,S,B)* reflects the sigmoidal transfer function for the total input *I* given within the curly braces:

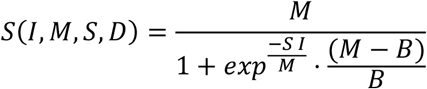

where *M* reflects the maximum firing rate, *S* the slope of the sigmoid, and *D* the spontaneous firing rate (i.e., baseline firing rate in the absence of input). Many of the values of these parameters can be ascertained from empirical estimates available from online databases (see supplementary table I). The model includes finite transmission delays using delayed values of *R*, i.e., the delayed input from the *j*^*th*^ to the *i*^*th*^ population is given by:

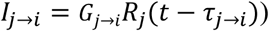

where *τ*_*j→i*_ reflects the finite time delay. Each state receives stochastic innovations added to the deterministic equations (given above). Delays were discretized and rounded to the nearest integration step size. Stochastic inputs were given by rescaling the variance of the noise to match the square root of the integration step *h* (i.e., *dW*_*t*_ *= W*_*t+h*_ *− W*_*t*_*∼N*(0, *h*), where *W*_*t*_ is a Wiener process, and *N* refers to the normal distribution. The system of equations was then integrated using an Euler-Maruyama scheme with fixed step size of 0.5 ms.

#### 6.5.5 Supplementary Methods V– Construction of Bifurcation Diagrams

Stability analysis was performed on a deterministic version of the model. This was achieved by setting the input constant and equal to the mean of the stochastic process. Initial conditions were found by running the stochastic model for 30s simulation time (by which models are at steady state) and taking the mean activity over states for the last 2s. To find equilibria, this deterministic model was simulated again for 30s, and inflection points in the states were identified by finding points at which the derivative was approximately zero. Unique equilibria (again determined within a set tolerance to the difference between equilibria) were then plot against the control parameter to construct bifurcation diagrams. We assessed the stability of the equilibria by computing eigenvalues λ of the (delayed adjusted) Jacobian at each fixed point (David et al. 2006). For the four state model of the motor cortex, this yields 4 (potentially complex-valued) eigenvalues for each value of the control parameter.

### 6.6 Supplementary Information 1 – Full ethics statements

The following ethics statements appear in their original, unmodified state supplied alongside the data repository.

#### Cued Finger Movements

“All patients participated in a purely voluntary manner, after providing informed written consent, under experimental protocols approved by the Institutional Review Board of the University of Washington (#12193). All patient data was anonymized according to IRB protocol, in accordance with HIPAA mandate. These data originally appeared in the manuscript “Human Motor Cortical Activity Is Selectively Phase-Entrained on Underlying Rhythms” published in PLoS Computational Biology in 2012 (Miller et al. 2012).”

#### Movement Imagery

“All patients participated in a purely voluntary manner, after providing informed written consent, under experimental protocols approved by the Institutional Review Board of the University of Washington (#12193). Portions of these data originally appeared in the manuscript “Cortical activity during motor execution, motor imagery, and imagery-based online feedback” published in PNAS in 2010 (Miller et al. 2010). Portions of these patient data was anonymized according to IRB protocol, in accordance with HIPAA mandate. It was made available through the library described in “A Library of Human Electrocorticographic Data and Analyses” by Kai Miller (Miller 2019), freely available at https://searchworks.stanford.edu/view/zk881ps0522.“

#### Basic Motor

“Ethics statement: All patients participated in a purely voluntary manner, after providing informed written consent, under experimental protocols approved by the Institutional Review Board of the University of Washington (#12193). All patient data was anonymized according to IRB protocol, in accordance with HIPAA mandate. It was made available through the library described in “A Library of Human Electrocorticographic Data and Analyses” by Kai Miller (Miller 2019), freely available at https://searchworks.stanford.edu/view/zk881ps0522. All patient data was anonymized according to IRB protocol, in accordance with HIPAA mandate. These data originally appeared in the manuscript “Spectral Changes in Cortical Surface Potentials during Motor Movement” published in Journal of Neuroscience in 2007 (Miller et al. 2007).”

### 6.7 Supplementary Table I – Prior Model Parameters

Where possible we derived prior estimates from empirical sources available from either the Allen Brain Atlas, or Neuroelectro.org. Estimates derived from human cells were preferred, but when not available, estimates in animals were also used. Estimates of prior precision (i.e., inverse variance) were obtained by looking at the variance in independently reported measurements.

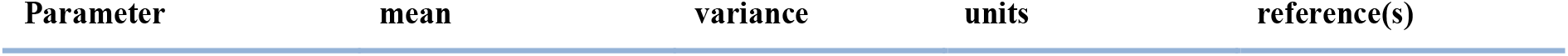
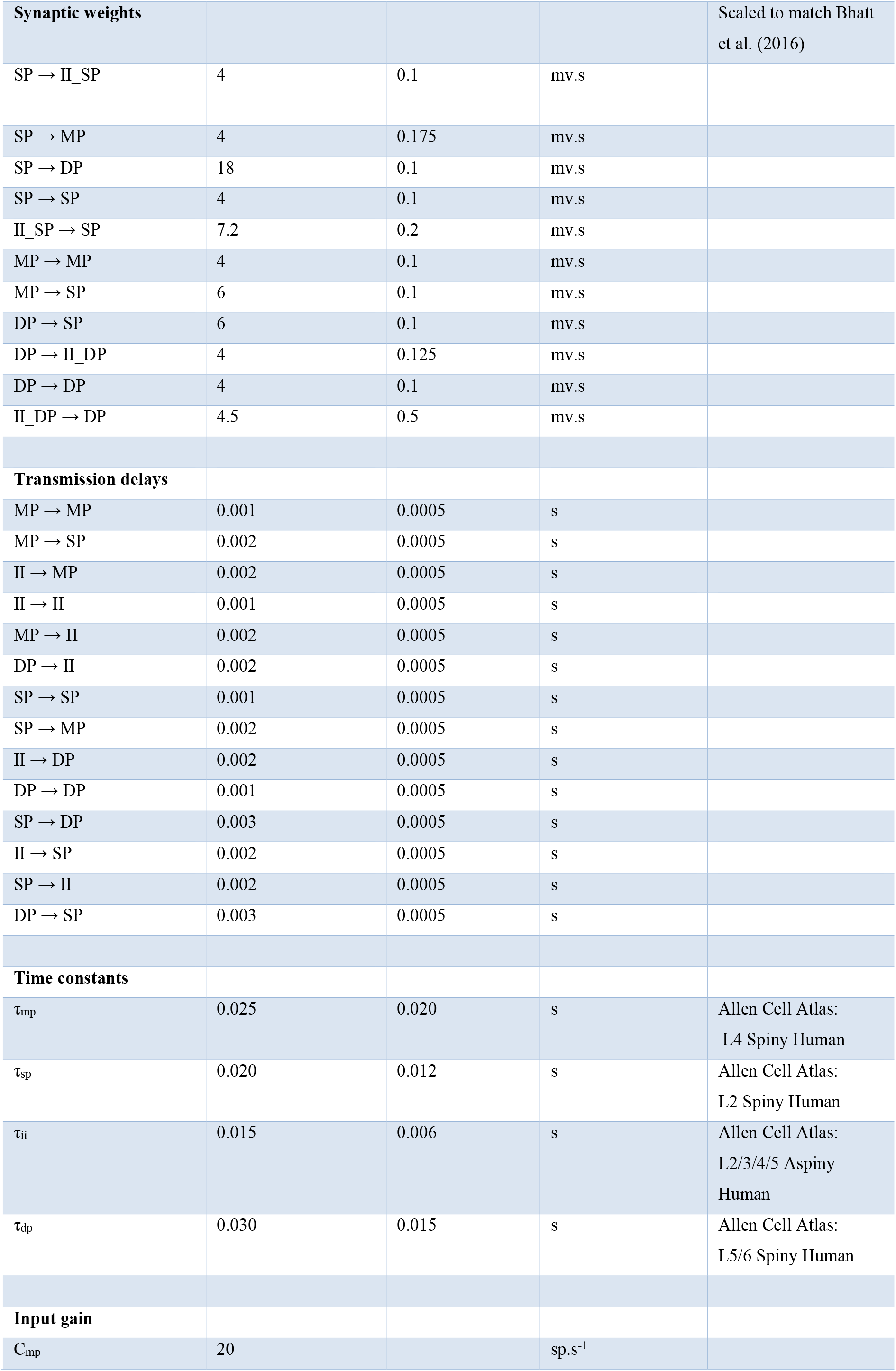
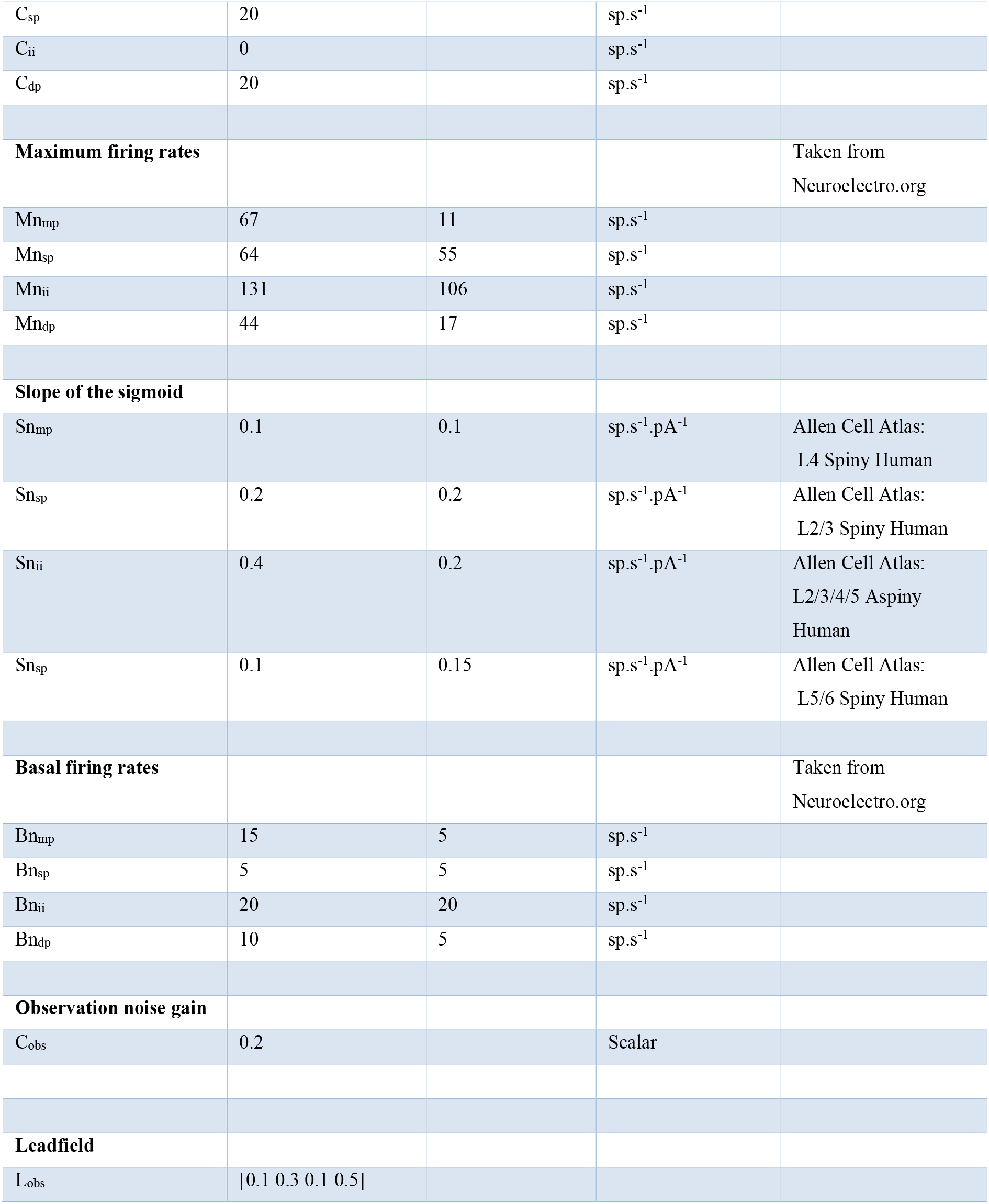

